# Gaining insights from RNA-Seq data using iDEP

**DOI:** 10.1101/148411

**Authors:** Steven Xijin Ge, Eun Wo Son

## Abstract

The analysis and interpretation of the RNA-Seq data can be time-consuming and challenging. We aim to streamline the bioinformatic analyses of gene-level data by developing a user-friendly web application for exploratory data analysis, differential expression, and pathway analysis. iDEP (integrated Differential Expression and Pathway analysis) seamlessly connects 63 R/Bioconductor packages, 208 annotation databases for plant and animal species, and 2 web services. The workflow can be reproduced by downloading customized R code and related files. As demonstrated by two examples, iDEP (http://ge-lab.org/idep/) democratizes access to bioinformatics resources and empowers biologists to easily gain actionable insights from transcriptomic data.

## Background

RNA sequencing (RNA-Seq) [1] is widely used for transcriptomic profiling. At increasingly reduced cost, library construction and sequencing can often be easily carried out following standard protocols. For many researchers, especially those without bioinformatics experience, the bottleneck to fully leverage the power of the technique is how to translate expression profiles into actionable insights. A typical analytic workflow involves many steps, each requiring several software packages. It can be time-consuming to learn, tune and connect these packages correctly. Another hurdle is the scattered annotation databases which use diverse types of gene IDs. To mitigate these issues, we aim to develop an application that can greatly reduce the time and effort required for researchers to analyze RNA-Seq data.

RNA-Seq data analysis often starts with quality control, pre-processing, mapping and summarizing of raw sequencing reads. We assume these steps were completed, using either the traditional Tuxedo Suite [2, 3] or alternatives such as the faster, alignment-free quantification methods [4, 5]. These tools can be used in stand-alone mood or through platforms like GenePattern [6] or web-based solutions such as Galaxy [7] and iPlant/CyVerse [8].

After read mapping, we often obtain a matrix of gene-level read counts or normalized expression levels (Fragments Per Kilobase Million, or FPKM). For such tabular data, like DNA microarray data, R is a powerful tool for visualization, exploratory data analysis (EDA), and statistical analysis. In addition, many dedicated R and Bioconductor [9] packages have been developed to identify differentially expressed genes (DEGs) and altered pathways. Some of the packages, such as DESeq2 [10], have been widely used. But it can be time-consuming, or even out of reach for researchers without coding experience.

Several web applications have been developed to analyze summarized expression data (Table 1). START App (Shiny Transcriptome Analysis Resource Tool) is a Shiny app that performs hierarchical clustering, principal component analysis (PCA), gene-level boxplots, and differential gene expression [11]. Another similar tool, Degust (http://degust.erc.monash.edu/) can perform differential expression analysis using EdgeR [12] or limma-voom [13] and interactively plot the results. Other tools include Sleuth [14] and ShinyNGS (https://github.com/pinin4fjords/shinyngs). Non-Shiny applications were also developed to take advantage of the R code base. This includes DEIVA [15] and VisRseq [16]. Beyond differential expression, several tools incorporate some capacity of pathway analysis. For quantified expression data, ASAP (Automated Single-cell Analysis Pipeline) [17] can carry out normalization, filtering, clustering, and enrichment analysis based on Gene Ontology (GO)[18] and KEGG [19] databases. With EXPath Tool [20], users can perform pathway search, GO enrichment and co-expression analysis. The development of these tools in the last few years facilitated the interpretation of RNA-Seq data.

**Table 1.**
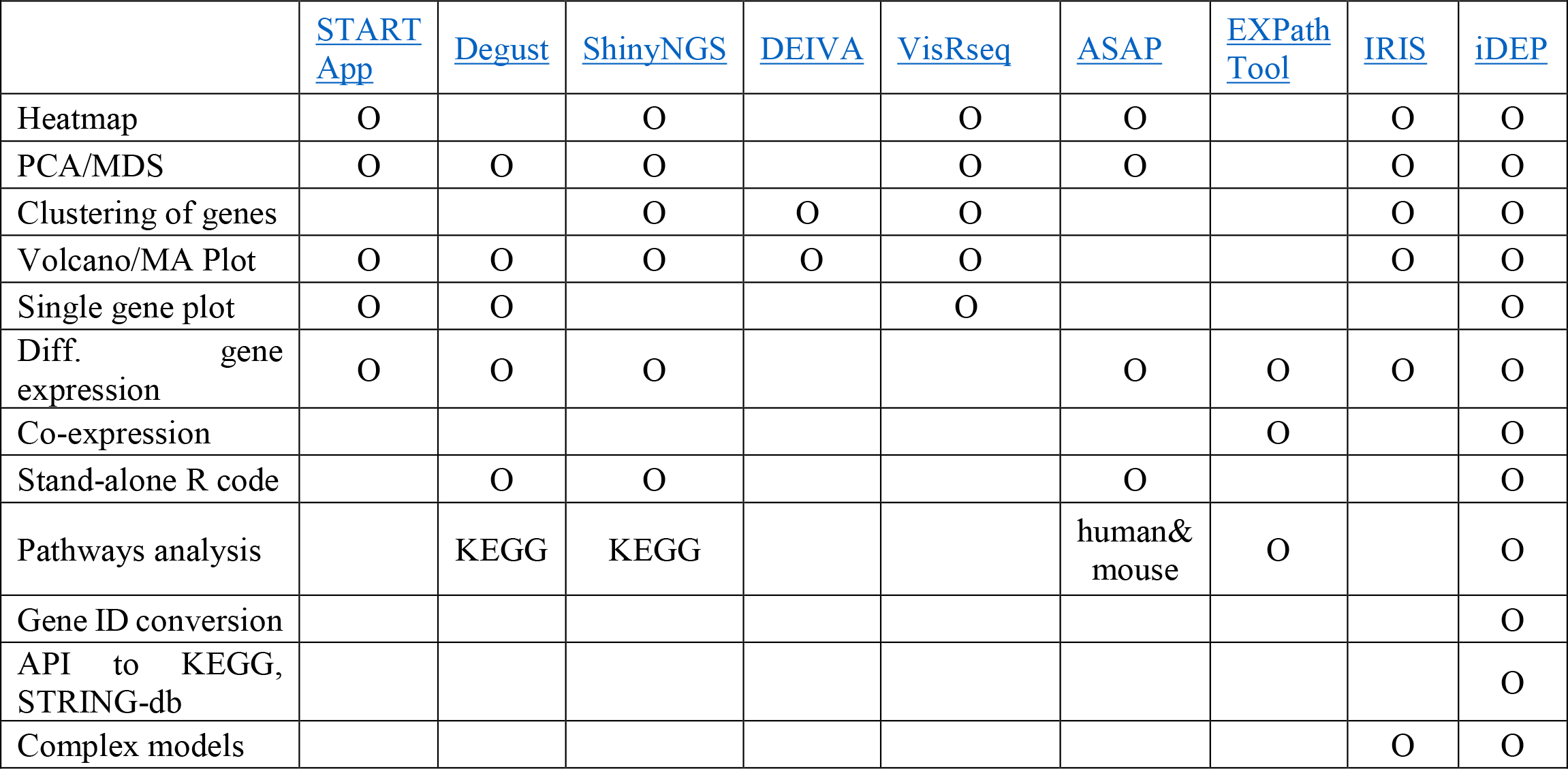
Comparison of applications for analyzing RNA-Seq.

|  | START App | Degust | ShinyNGS | DEIVA | VisRseq | ASAP | EXPath Tool | IRIS | iDEP |
| --- | --- | --- | --- | --- | --- | --- | --- | --- | --- |
| Heatmap | O |  | O |  | O | O |  | O | O |
| PCA/MDS | O | O | O |  | O | O |  | O | O |
| Clustering of genes |  |  | O | O | O |  |  | O | O |
| Volcano/MA Plot | O | O | O | O | O |  |  | O | O |
| Single gene plot | O | O |  |  | O |  |  |  | O |
| Diff. gene expression | O | O | O |  |  | O | O | O | O |
| Co-expression |  |  |  |  |  |  | O |  | O |
| Stand-alone R code |  | O | O |  |  | O |  |  | O |
| Pathways analysis |  | KEGG | KEGG |  |  | human& mouse | O |  | O |
| Gene ID conversion |  |  |  |  |  |  |  |  | O |
| API to KEGG, STRING-db |  |  |  |  |  |  |  |  | O |
| Complex models |  |  |  |  |  |  |  | O | O |

In this study, we seek to develop a web application with substantially enhanced functionality with (1) automatic gene ID conversion with broad coverage, (2) comprehensive gene annotation and pathway database for both plant and animals, (3) several methods for in-depth EDA and pathway analysis, (4) access to web services such as KEGG [19] and STRING-db [21] via application programming interface (API), and (5) improved reproducibility by generating R scripts for stand-alone analysis.

Leveraging many existing R packages (see Figure 1) and the efficiency of the Shiny framework, we developed iDEP (integrated Differential Expression and Pathway analysis) to enable users to easily formulate new hypotheses from transcriptomic datasets. Taking advantage of the massive amount of gene annotation information in Ensembl [22, 23] and Ensembl Plants [24], as well as pathway databases compiled by our group [25, 26] and others [19, 27, 28], we were able to build a large database to support in-depth pathway analyses.

**Figure 1.**
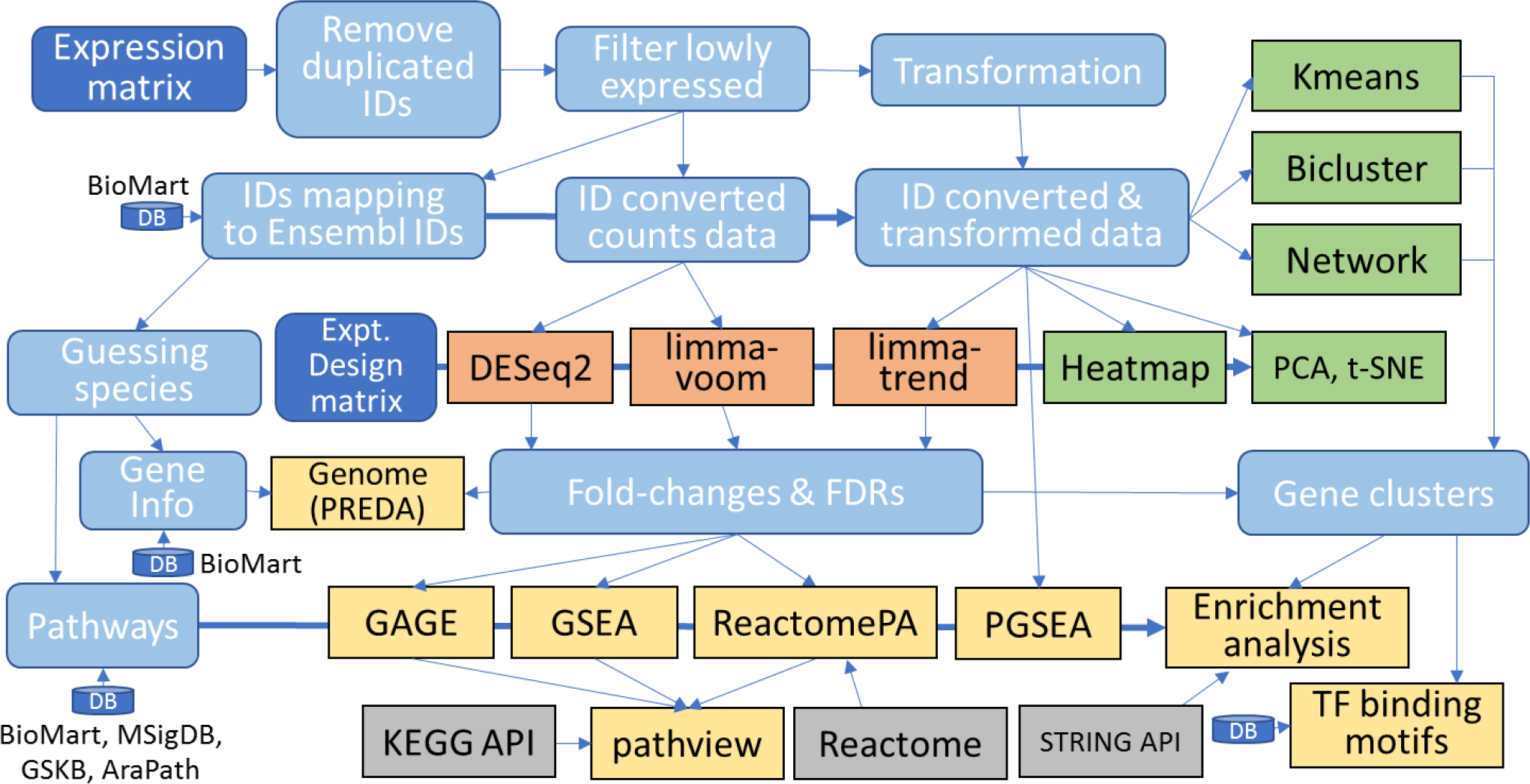
iDEP functional modules.

We used iDEP to analyze two example datasets and generate all the figures and tables in this paper except Table 1 and Figure 1. We first used it to extensively analyzed a simple RNA-Seq dataset involving small interfering RNA (siRNA)-mediated Hoxa1 knockdown in human lung fibroblasts [3]. We identified the down-regulation of cell-cycle genes, in agreement with previous studies. Our analyses also reveal the possible roles of E2F1 and its target genes, including microRNAs, in blocking G_1_/S transition, and the upregulation of genes related to cytokines, lysosome, and neuronal parts. The second dataset was derived from an experiment with a factorial design to study the effect of ionizing radiation (IR) on mouse B cells with and without functional TRP53 [29]. In addition to correctly identifying p53 pathway and the enrichment of p53 target genes, we also found the p53-independent effects, including the regulation of ribosome biogenesis and non-coding RNA metabolism, and activation of c-MYC. These examples show that users can gain insights into both molecular pathways and gene regulatory mechanisms.

## Results

We developed an easy-to-use web application that encompasses many useful R and Bioconductor packages, vast annotation databases, and related web services. The input is a gene-level expression matrix obtained from RNA-seq, DNA microarray, or other platforms. Main functionalities include (1) pre-processing, (2) EDA, (3) differential expression, and (4) pathway analysis and visualization.

We tried to develop an intuitive, graphical, and robust tool so that researchers without bioinformatics experience can routinely and quickly translate expression data into novel hypotheses. We also wanted to make an open system where users can download intermediate results so that other tools can be used. Also, users can upload custom pathway databases for unannotated species. For experienced bioinformaticians, it can serve as a tool for preliminary analysis as it circumcises the need for many tedious tasks such as converting gene IDs and downloading software packages and annotations. These users can also download customized R scripts and related data files so that the analysis can be reproduced and extended.

### iDEP Design

Figure 1 outlines the iDEP workflow. Expression matrix is first filtered, transformed and converted to Ensemble gene IDs, which are used internally to identify genes. The pre-processed data is then used for EDA, such as K-means clustering, hierarchical clustering, principal component analysis (PCA), t-SNE[30]. Gene clusters identified by K-means are analyzed by enrichment analysis based on a large gene annotation and pathway database. The identification of DEGs is done with either the *limma* [31] or DESeq2 [10] packages. This is also followed by enrichment analysis on the DEGs. The fold-change values are then used in pathway analysis using several methods.

To enable gene ID conversion, we downloaded all available gene ID mappings for 208 species from Ensembl [22, 23] (Table S1 in Supp. File 1), including 97 from Ensembl (vertebrates, release 91), 45 from Ensembl Plants (release 37) [24], and 66 from Ensembl Metazoa (release 37). The final mapping table for the current iDEP v0.72 release consists of 135,832,098 rows, mapping various gene IDs (Table S2 in Supp. File 1) into Ensembl. For example, 67 types of human gene IDs can be converted to Ensembl gene IDs.

Besides common ID like gene symbol, Entrez, Refseq, UCSC, UniGene, and Interpro IDs, the 67 kinds of human gene IDs also include probe IDs for popular DNA microarray platforms, making it possible to re-analyze thousands of microarray datasets available at public repositories.

In the pre-processing stage, gene IDs are first compared to all gene IDs in the database for 208 organisms. This enables automatic ID conversion and species identification. Genes expressed at very low levels are removed and data are transformed as needed using one of several methods. iDEP enforces log-transformation when a highly skewed distribution is detected. This type of mechanisms can help avoid issues in downstream analyses. The pre-processing stage also generates diagnostic and summary plots to guide users to make their choices.

EDA enables the users to explore variations and patterns in the dataset as a whole [32]. The main methods include hierarchical clustering with heatmap, k-means clustering, and PCA. Enrichment analysis of genes derived from k-means clustering is conducted to gain insights into the functions of co-expressed genes. Initial attempts of pathway analysis are carried out using the PCA loadings on each gene. This can tell us the biological processes underlying each direction of expression change defined by the principal components.

Differential expression analysis relies on two Bioconductor packages, *limma* [31] and DESeq2 [10]. These two packages can meet the needs for most studies, including those involving multiple biological samples and factorial design. Normalized expression data is analyzed using *limma*. Read counts data can be analyzed using three methods, namely limma-trend [13], limma-voom [13, 33], and DESeq2. Other methods such as edgeR [12] may be incorporated in the future.

For simple study designs, iDEP runs differential gene expression analysis on all pairs of sample groups, which are defined by parsing sample names. For complex studies, users can upload a file with experiment design information and then build statistical models that can involve up to 6 factors. This also enables users to control for batch effects or dealing with paired samples.

Fold-change values for all genes returned by *limma* or DESeq2 are used in pathway analysis using GSEA [34], PAGE [35, 36], GAGE[37] or ReactomePA [38]. Taking advantage of centralized annotation databases for 208 species at Ensembl, Ensembl Plants, and Ensembl Metazoa, we downloaded not only GO functional categorizations, but also promoter sequences for defining transcription factor (TF) binding motifs for most species. Metabolic pathways were downloaded directly from KEGG [19] for 117 species (Table S1 in Supp. File 1). Also, we incorporated Pathview package [39] to show gene expression on KEGG pathway diagrams downloaded via API. In addition, we also included many species-specific pathway knowledgebases, such as Reactome [28, 38], GeneSetDB[40] and MSigDB [27] for human, GSKB for mouse [25], and araPath for Arabidopsis [26]. These databases contain diverse types of gene sets, ranging from TF and microRNA target genes to manually curated lists of published DEGs. For the human genome, we collected 72,394 gene sets (Table 2). Such comprehensive databases enable in-depth analysis of expression data from different perspectives.

**Table 2.**
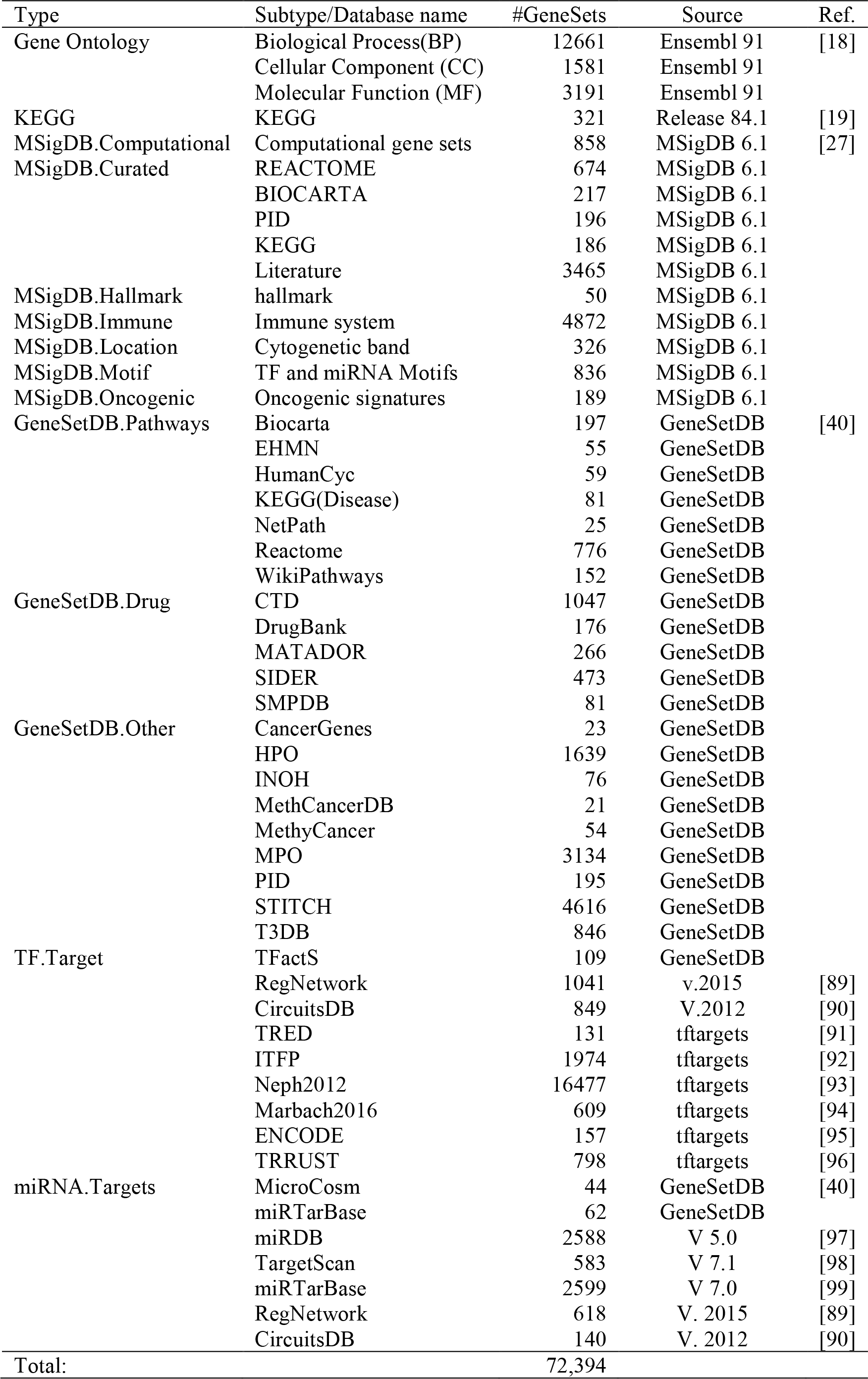
Gene set databases collected for enrichment analysis in human.

iDEP also enables users to retrieve protein-protein interaction (PPI) networks among top DEGs via an API access to STRING [21]. These networks can be rendered both as static images and as richly annotated, interactive graphs on the STRING website. The API access also provides enrichment analysis (GO, KEGG, and protein domains) for 115 archaeal, 1678 bacterial, and 238 eukaryotic species, thus greatly expanding the species coverage of iDEP.

Based on their chromosomal location obtained from Ensembl, we visualize fold-changes of genes on all the chromosomes as an interactive graph based on Plotly. iDEP can also use the PREDA package [41] to detect chromosomal regions overrepresented with up- or down-regulated genes. This is useful for studies such as cancer that might involve chromosomal deletion or amplification.

For larger datasets, users can use bi-clustering algorithms to identify genes with correlated expression among a subset of samples, using the 8 methods implemented in 3 Bioconductor packages biclust [42], QUBIC [43], and runibic [44]. Gene co-expression networks can also be constructed with the WGCNA package [45]. Enrichment analysis is routinely conducted on gene clusters derived from these methods.

To ensure the ease of deployment, we used docker containers to configure the web server. This technology also enables us to easily clone the service many times to take advantage of the multiple cores. Load balanced with Nginx, our web server can handle hundreds of concurrent users. The source code for iDEP and our server configuration files are available at our Github repository (https://github.com/iDEP-SDSU/idep).

### Use case 1: a simple experiment on Hoxa1 knockdown

We first analyzed a simple dataset related to Hoxa1 knockdown by siRNA in human lung fibroblasts[3]. With 3 replicates for each of the two biological samples, this RNA-Seq dataset was used as example data for Cuffdiff2 [3]. Available as Supplementary File 2, the read count data was previously used in a tutorial for pathway analysis [46].

#### Pre-processing and EDA

After uploading the read count data, iDEP correctly recognized Homo sapiens as the likely species with the most genes IDs matched. After ID conversion and the default filter (0.5 counts per million, or CPM, in at least one sample), 13,819 genes left. A bar plot of total read counts per library is generated (Figure 2A), showing some small variation in library sizes. We chose the regularized log (rlog) transformation implemented in the DESeq2 package, as it effectively reduces mean-dependent variance (Figure S1 in Supp. File 3). Distribution of the transformed data is shown in Figure 2B-C. Variation among replicates is small (Figure 2D).

**Figure 2.**
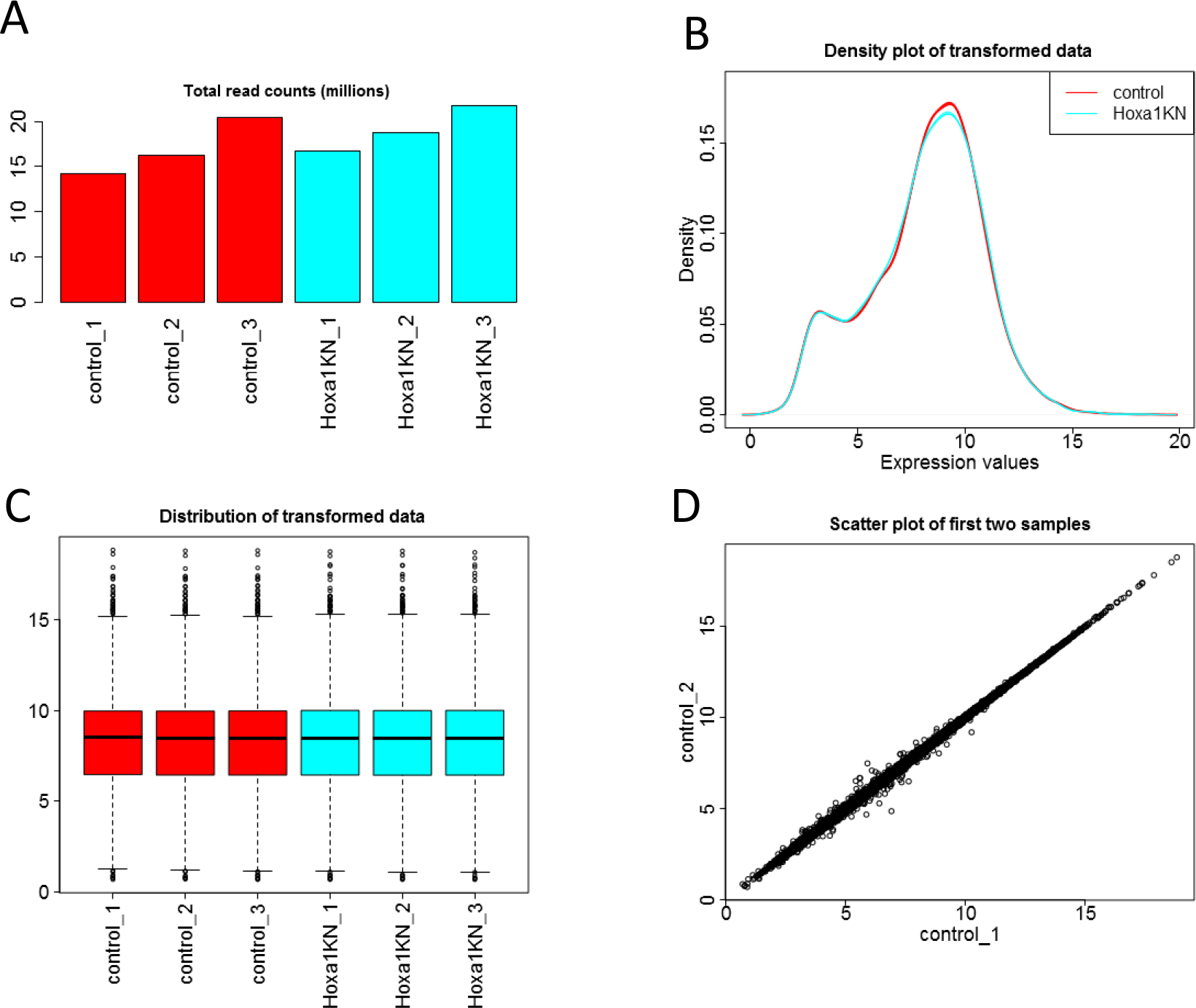
Diagnostic plots for read-counts data.A) Total read-counts per library. B) Distribution of transformed data using a density plot. C) Boxplot of transformed data. D) Scatter plot of the first two samples.

iDEP also enables users to examine the expression level of one or more genes. Using “Hoxa” as a keyword, we obtain Figure 3A, which shows that Hoxa1 expression level is reduced, but not abolished, in response to siRNA-mediated knockdown of Hoxa1. Noticeably, expression of other family members, especially Hoxa2, 4, and 5, also decrease. As these genes have similar mRNA sequences, it is unclear whether this is caused by off-target effects of the siRNA or ambiguous mapping of RNA-Seq reads. Figure 3B, obtained by using “E2F” as a keyword, shows the down-regulation of E2F1.

**Figure 3.**
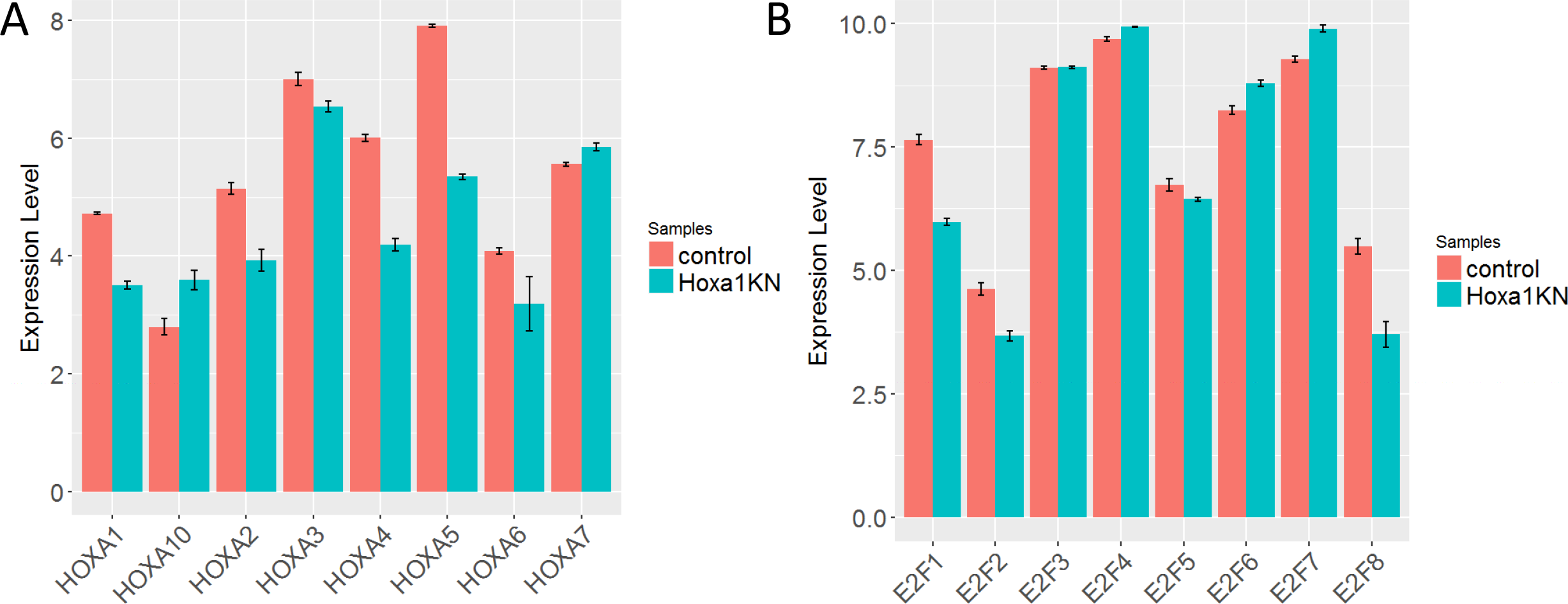
Expression patterns of (A) Hoxa and (B) E2f gene families.

For clustering analysis, we rank genes by their standard deviation across all samples. The result of hierarchical clustering of the top 1000 genes is shown in Figure 4A, which suggests that Hoxa1 knockdown in lung fibroblast cells induce a drastic change in the expression of hundreds of genes. Variations among technical replicates are minimal. These observations can also be confirmed by the correlation matrix (Supp. Figure 2) and k-means clustering (Supp. Figure 3).

**Figure 4.**
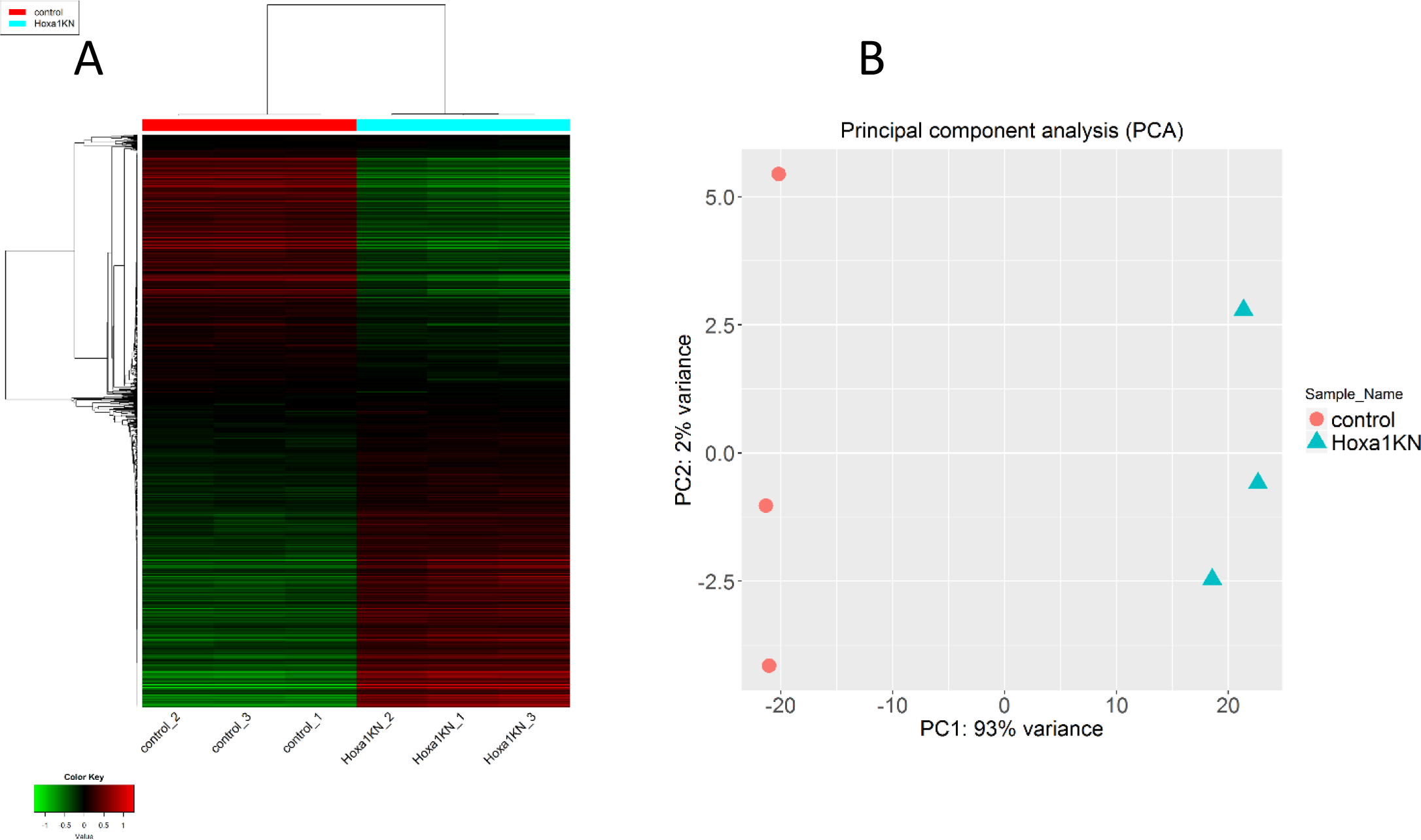
Hierarchical clustering (A) and PCA analyses(B) indicate the substantial difference in thousands of genes induced by Hoxa knockdown. There is little variation among replicates.

PCA plot using the first and second principal components is shown in Figure 4B. There is a clear difference between the Hoxa1 knockdown and the control samples, along the first principal component that explains 93% of the variance. Plot using multidimensional scaling (MDS), and t-SNE [30] also show a similar distribution of the samples (Supp. Figure 4). We can choose to conduct pathway analysis using PGSEA [35, 36] by treating the loadings of the principal components as expression values. As suggested by Supp. Figure 5, the first two components are related to cell cycle regulation.

**Figure 5.**
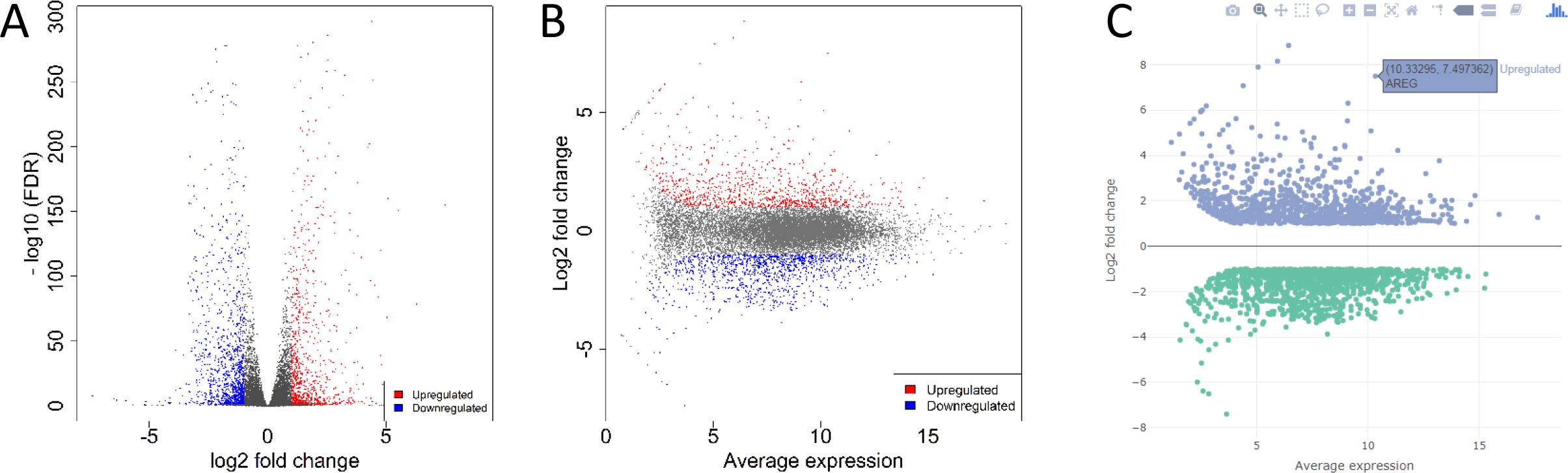
Summary plots for differential expression analysis using DESeq2. A) Volcano plot, B) M-A plot, and C) Interactive MA plot.

#### Differentially expressed genes (DEGs)

With the DESeq2 method, we identified 907 upregulated and 1097 downregulated genes (see Supp. Table 3) using a threshold of false discovery rate (FDR) < 0.1 and fold-change >2. The volcano plot (Figure 5A) and the M-A plot (Figure 5B) show that Hoxa1 knockdown leads to a massive transcriptomic response. Plotly-based interactive versions of these plots are also available, where users can zoom in and mouse over to see individual genes (Figure 5C). A quick scan at the top genes ranked by the absolute values of fold-change (FCs) tells us that Hoxa1 knockdown induces cytokines (IL1B, IL24).

**Table 3.** Enriched GO terms in up and down-regulated genes.

| Direction | adj.Pval | nGenes | Pathways |
| --- | --- | --- | --- |
| Down<br>Regulated | 4.50E-46 | 259 | Cell cycle |
|  | 1.70E-42 | 207 | Cell cycle process |
|  | 5.60E-38 | 169 | Mitotic cell cycle |
|  | 2.80E-35 | 92 | Chromosome segregation |
|  | 2.00E-33 | 145 | Mitotic cell cycle process |
|  | 5.20E-33 | 72 | Sister chromatid segregation |
|  | 5.20E-33 | 118 | Cell division |
|  | 2.20E-30 | 80 | Nuclear chromosome segregation |
|  | 5.60E-26 | 86 | Nuclear division |
|  | 1.20E-24 | 350 | Organelle organization |
|  | 3.50E-24 | 65 | Mitotic nuclear division |
|  | 5.80E-24 | 88 | Organelle fission |
|  | 1.70E-23 | 159 | Cytoskeleton organization |
|  | 2.60E-21 | 98 | Cell cycle phase transition |
|  | 4.00E-21 | 45 | Mitotic sister chromatid segregation |
| Up<br>Regulated | 3.40E-25 | 258 | Cell surface receptor signaling pathway |
|  | 8.20E-23 | 171 | Regulation of cell proliferation |
|  | 7.40E-22 | 195 | Cell proliferation |
|  | 3.80E-21 | 265 | Regulation of signaling |
|  | 1.20E-20 | 259 | Response to organic substance |
|  | 1.20E-20 | 260 | Regulation of cell communication |
|  | 1.20E-19 | 165 | Locomotion |
|  | 1.30E-19 | 239 | Regulation of signal transduction |
|  | 1.00E-18 | 332 | System development |
|  | 3.70E-17 | 100 | Regulation of cellular component movement |
|  | 8.20E-17 | 291 | Regulation of response to stimulus |
|  | 1.20E-16 | 143 | Response to endogenous stimulus |
|  | 1.70E-16 | 319 | Response to chemical |
|  | 1.80E-16 | 211 | Cellular response to organic substance |
|  | 2.10E-16 | 131 | Cell migration |

The up and down-regulated genes are then subjected to enrichment analysis based on the hypergeometric distribution. Many different types of genes sets listed in Table 2 can be used to test various hypotheses. The GO Biological Process terms enriched in DEGs are shown in Table 3. Upregulated genes are related to regulation of cell proliferation, locomotion, and response to endogenous stimuli. This is perhaps the cell’s response to injected siRNAs. The downregulated genes are significantly enriched with cell cycle-related genes (FDR < 2.6x10^−47^). The effect of Hoxa1 knockdown on cell cycle was reported and experimentally confirmed in the original study [3]. Cell cycle analysis revealed that loss of Hoxa1 leads to a block in G_1_ phase [3].

As many GO terms are related or redundant (*i.e*., cell cycle and cell cycle process), we provide two plots to summarize such correlation [47]. We first measure the distance among the terms by the percentage of overlapped genes. Then this distance is used to construct a hierarchical tree (Figure 6A) and a network of GO terms (Figure 6B). Both plots show that the enriched terms are distinct in the two gene lists. The down-regulated genes are overwhelmingly involved in cell cycle. The upregulated genes are related to 4 related themes: cell proliferation, signaling, response to organic substance, and cell migration, possibly in reaction to the injected siRNAs.

**Figure 6.**
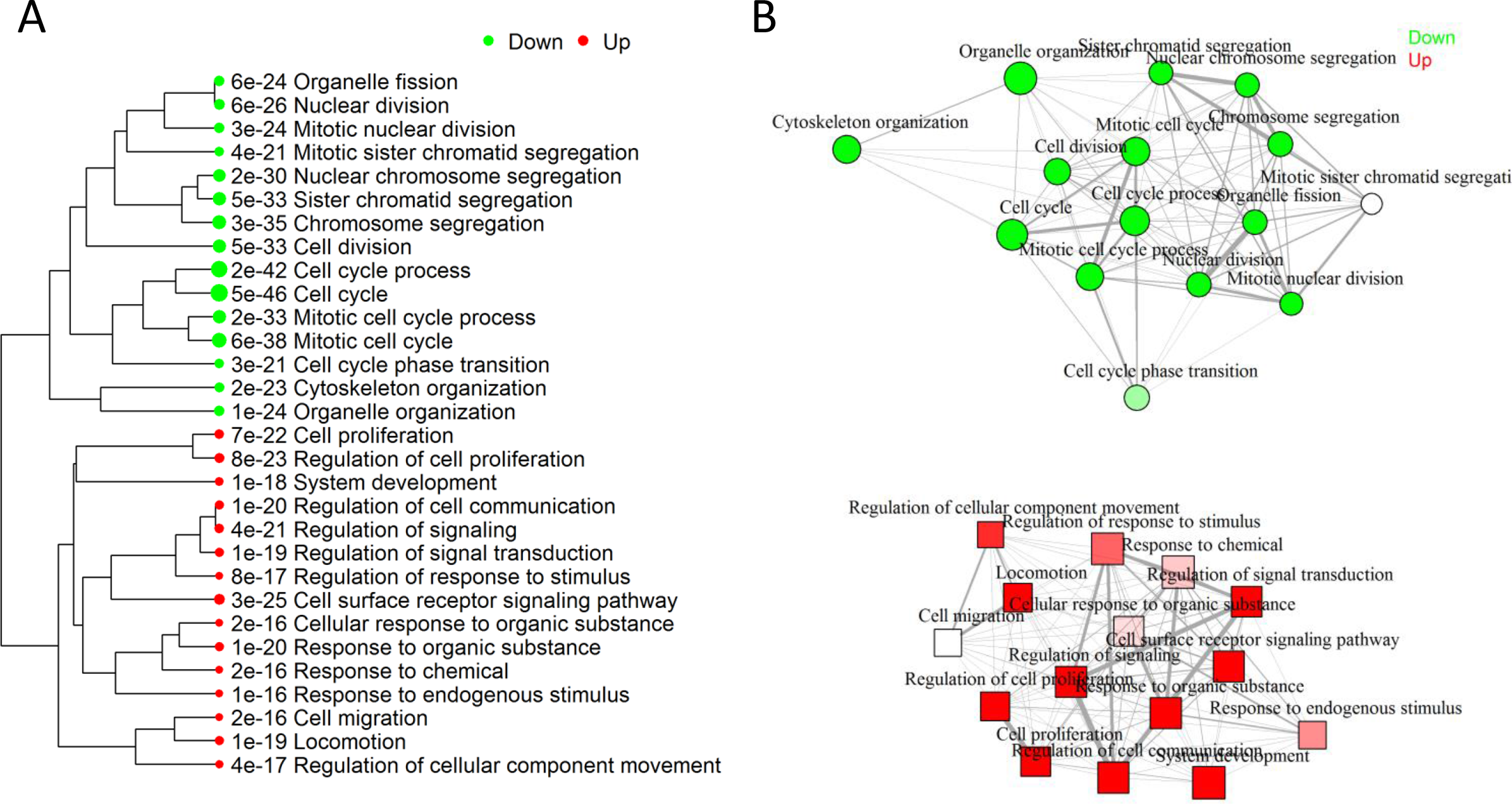
Visualization of the relationship among enriched GO categories using (A) tree and (b) network.

Choosing GO cellular component, we find that Hoxa1 knockdown suppresses genes that code for the spindle, cytoskeleton and chromosomal parts (Supp. Figure 6). As Hoxa1 knockdown blocks G_1_/S transition [3], a smaller number of cells are in the S (synthesis) phase, leading to the reduction of proteins related to the spindle and chromosomal parts. Hoxa1 knockdown also induces genes related to plasma membrane, neurons and synapses (Supp. Figure 6). This unexpected result is consistent with Hoxa1’s role in neuronal differentiation [48, 49]. Polymorphisms of this gene are associated with cerebellar volume in humans [50]. Hoxa1 may have different functions in various organs across developmental stages.

Choosing KEGG pathway, we confirm the overrepresentation of cell cycle-related genes in downregulated genes (Supp. Figure 7). For up-regulated genes, we detect cytokine-cytokine receptor interaction (CCRI) pathway (FDR<1.3×10^−10^). “MSigDB.Curated” gene sets contain pathways from various databases as well as published lists of DEGs from previous expression studies [27]. As shown in Supp. Figure 8, the most significant are oligodendrocyte differentiation and several cell-cycle related gene sets. As suggested by a meta-analysis of published gene lists [51], cell-cycle related expression signature is frequently triggered by diverse cellular perturbations [52]. We uncovered similarity of our expression signature with previously published ones.

**Figure 7.**
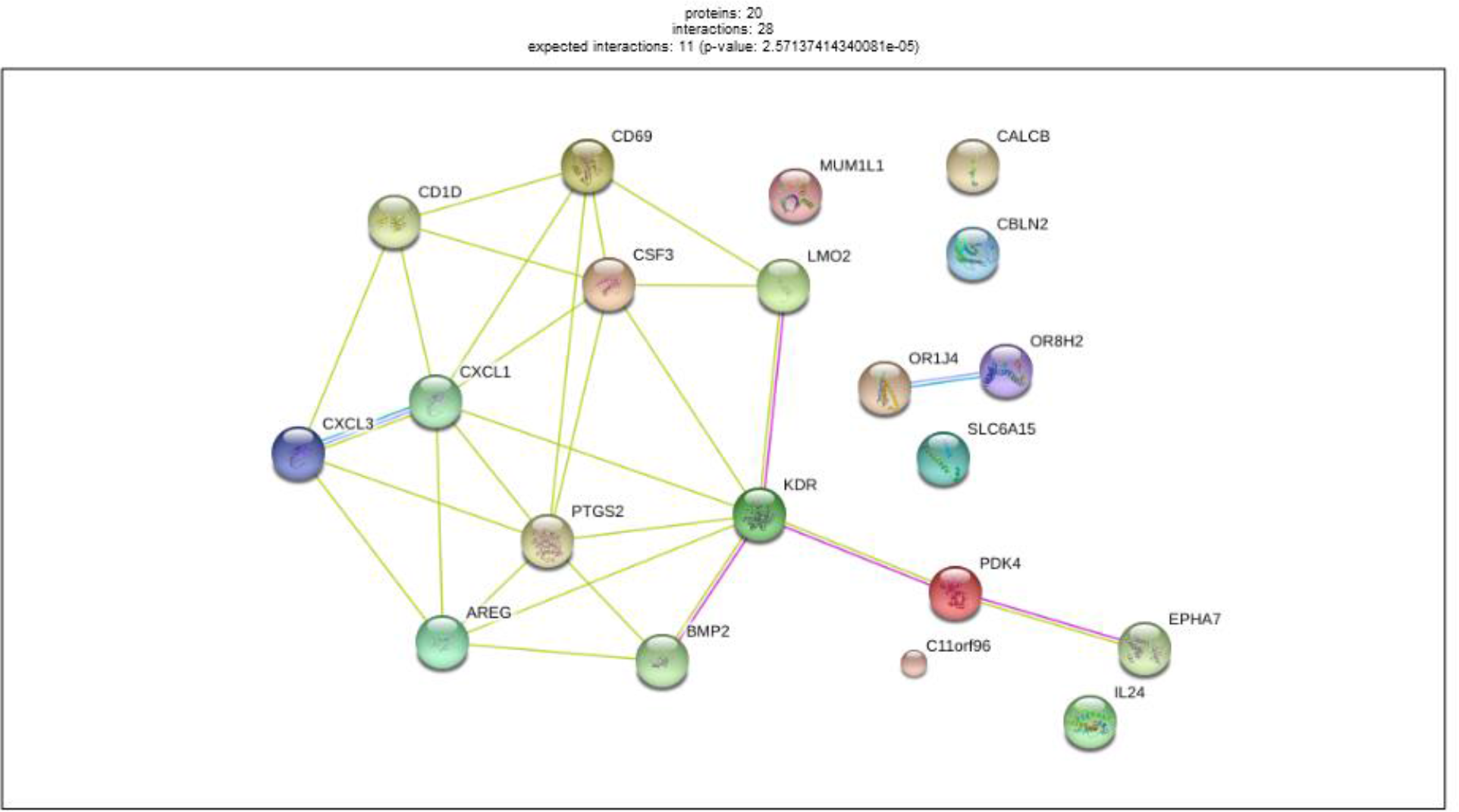
Protein-protein interactions (PPI) among top 20 up-regulated genes. This is retrieved via API access to STRING database. There is also an enrichment of PPIs compared with background. An interactive version of this network is also available through a link to the STRING website.

**Figure 8.**
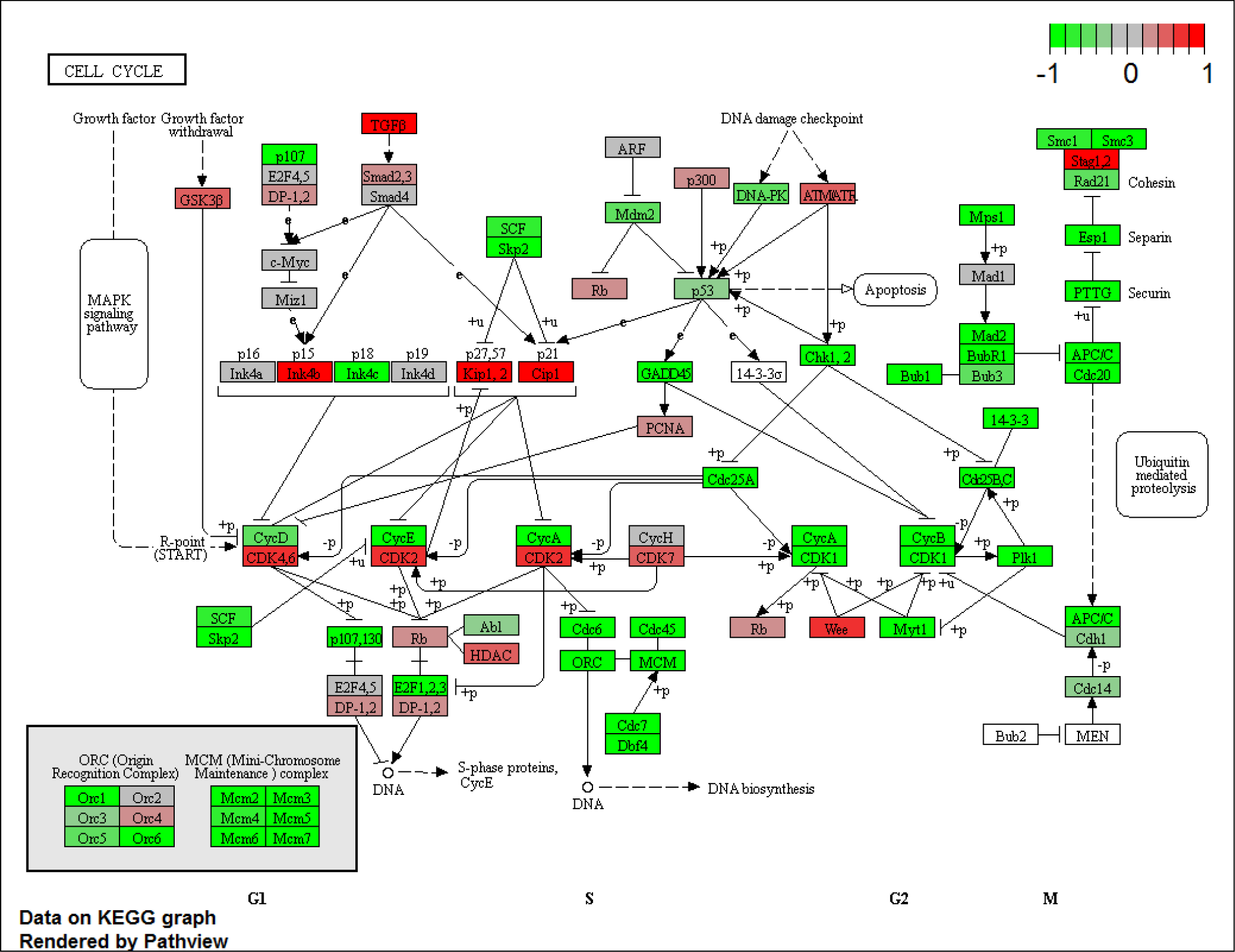
Expression profiles of cell-cycle related genes visualized on KEGG pathway diagram using the Pathview package. Red and green indicate genes induced or suppressed by Hoxa1 knockdown, respectively.

Using the STRINGdb Bioconductor package, iDEP can analyze the lists of DEGs via the STRING API [21] for enrichment analysis and the retrieval of PPI networks. The enrichment analysis led to similar results (Supp. Table 4) to those obtained using the internal iDEP gene sets. In addition, STRING detected that the Helix-loop-helix DNA-binding domain is overrepresented in proteins coded by the 907 upregulated genes, while the Tubulin/FtsZ family, GTPase domain is enriched in the 1097 down-regulated genes (Supp. Table S5). Figure 7 is the network of PPIs among the top 20 upregulated genes. The highly connected network includes chemokine ligands 1 and 3 (CXCL1 and CXCL3), as well as interleukin 24 (IL24), suggesting the immune response caused by injected siRNA. A link to an interactive version of this network on the STRING website is also available for access to rich annotation information including protein structures and literature.

**Table 4.** Enriched transcription factor(TF) binding motifs.

| Direction | adj.Pval | nGenes | Pathways |
| --- | --- | --- | --- |
| Down<br>Regulated | 9.80E-23 | 475 | Tftargets:TF Target SP1 |
|  | 9.80E-23 | 65 | Tftargets:TF Target E2F-4 |
|  | 1.10E-16 | 43 | TFactS E2F1 |
|  | 9.70E-15 | 37 | TRRUST:TF Target E2F1 |
|  | 1.20E-14 | 127 | RegNetwork:TF Target E2F4 |
|  | 1.00E-13 | 208 | RegNetwork:TF Target E2F1 |
|  | 1.30E-12 | 20 | TFactS E2F4 |
|  | 4.60E-09 | 154 | RegNetwork:TF Target NFYA |
|  | 1.80E-08 | 15 | TFactS E2F3 |
|  | 1.90E-08 | 33 | Tftargets:TF Target TEAD1 |
|  | 2.30E-08 | 74 | Tftargets:TF Target AP1 |
|  | 2.60E-08 | 14 | TFactS E2F2 |
|  | 3.30E-08 | 34 | Tftargets:TF Target TGIF1 |
|  | 4.50E-08 | 60 | Tftargets:TF Target ZNF219 |
|  | 5.00E-08 | 101 | Tftargets:TF Target HF1H3B |
| Up<br>Regulated | 4.90E-09 | 54 | Tftargets:TF Target NFKB |
|  | 4.90E-09 | 21 | TFactS FOXO3 |
|  | 7.30E-09 | 34 | Tftargets:TF Target NFKB1 |
|  | 7.30E-09 | 42 | TRRUST:TF Target NFKB1 |
|  | 4.40E-08 | 40 | TFactS CTNNB1 |
|  | 2.00E-07 | 22 | Tftargets:TF Target FOXJ1 |
|  | 4.00E-07 | 35 | Tftargets:TF Target POU3F2 |
|  | 4.20E-07 | 38 | TRRUST:TF Target RELA |
|  | 4.80E-07 | 28 | Tftargets:TF Target FOXO3 |
|  | 4.80E-07 | 50 | TRRUST:TF Target SP1 |
|  | 5.20E-07 | 25 | TRRUST:TF Target JUN |
|  | 6.10E-07 | 19 | TRRUST:TF Target EGR1 |
|  | 1.20E-06 | 39 | Tftargets:TF Target SP1 |
|  | 1.20E-06 | 25 | Tftargets:TF Target FOXJ3 |
|  | 1.20E-06 | 24 | Tftargets:TF Target FOXL1 |

iDEP can also reveal gene regulatory mechanisms. Using the TF target gene sets in enrichment analyses, we can obtain Table 4, which suggest that binding motifs for SP1 (FDR<9.80×10^−23^) and E2F factors (FDR<1.1×10^−16^) are enriched in the promoters of down-regulated genes. E2F factors are regulators of cell cycle [53]. E2F1 promotes G_1_/S transition [54] by regulation many genes, including itself. SP1 binding sites were identified in cell-cycle related genes such as Cyclin D1 (CCD1)[55]. SP1 is a G1 phase specific TF [56]. The interaction of E2F1 and SP1 proteins mediate cell cycle regulation [57]. The upregulated genes are enriched with target genes of NF-κB (FDR<4.9×10^−9^) and FOXO3(FDR<4.9×10^−9^), known to be regulators of the immune response [58, 59].

The Motif gene sets from MSigDB are derived from [60] and contain sets of genes sharing TF binding motifs in gene promoters and microRNA target motifs in 3’ untranslated regions (UTRs). Using this gene set, we again detect the enrichment of E2F motifs in promoters of downregulated genes (Supp.Table S10). We also detected overrepresentation of a “GCACTTT” motif in 3’ UTRs of upregulated genes. This motif is targeted by several microRNAs, namely miR-17-5P, miR-20a, miR-106a. Cloonan *et al*. showed that miR-17-5P targets more than 20 genes involved in the G_1_/S transition [61]. Trompeter *et al*. provided evidence that miR-17, miR-20a, and miR-106b enhance the activities of E2F factors to influence G_1_/S transition [62]. miR-106b resides in the intron of Mcm7 along the sense direction. Mcm7 is an E2F1 target gene that is also downregulated by Hoxa1 knockdown (see Fig. 8A). Petrocca *et al*. showed that E2F1 regulates miR-106b, which can conversely control E2F1 expression [63]. Thus, it is possible that Hoxa1 knockdown reduces E2F1 expression (see Figure 3B) and its target genes, including Mcm7, which hosts miR-106b. We can speculate that downregulated miR-106b, in turn, causes the increases in the expression of its target genes. Leveraging the comprehensive pathway databases, iDEP enables researchers to develop new hypotheses that could be further investigated.

For many species, predicted TF target genes are not available. As a solution, we downloaded promoter 300bp and 600bp sequences from ENSEMBL and scanned them with a large collection of TF binding motifs [64]. As shown in Table 5, the promoters of DEGs are overrepresented with many G-rich motifs bound by E2F and other factors such as TCFL5 and SP2. We compared the best possible scores for each TF and promoter pair and run t-tests to compare these scores. Further study is needed to validate this approach.

**Table 5.**
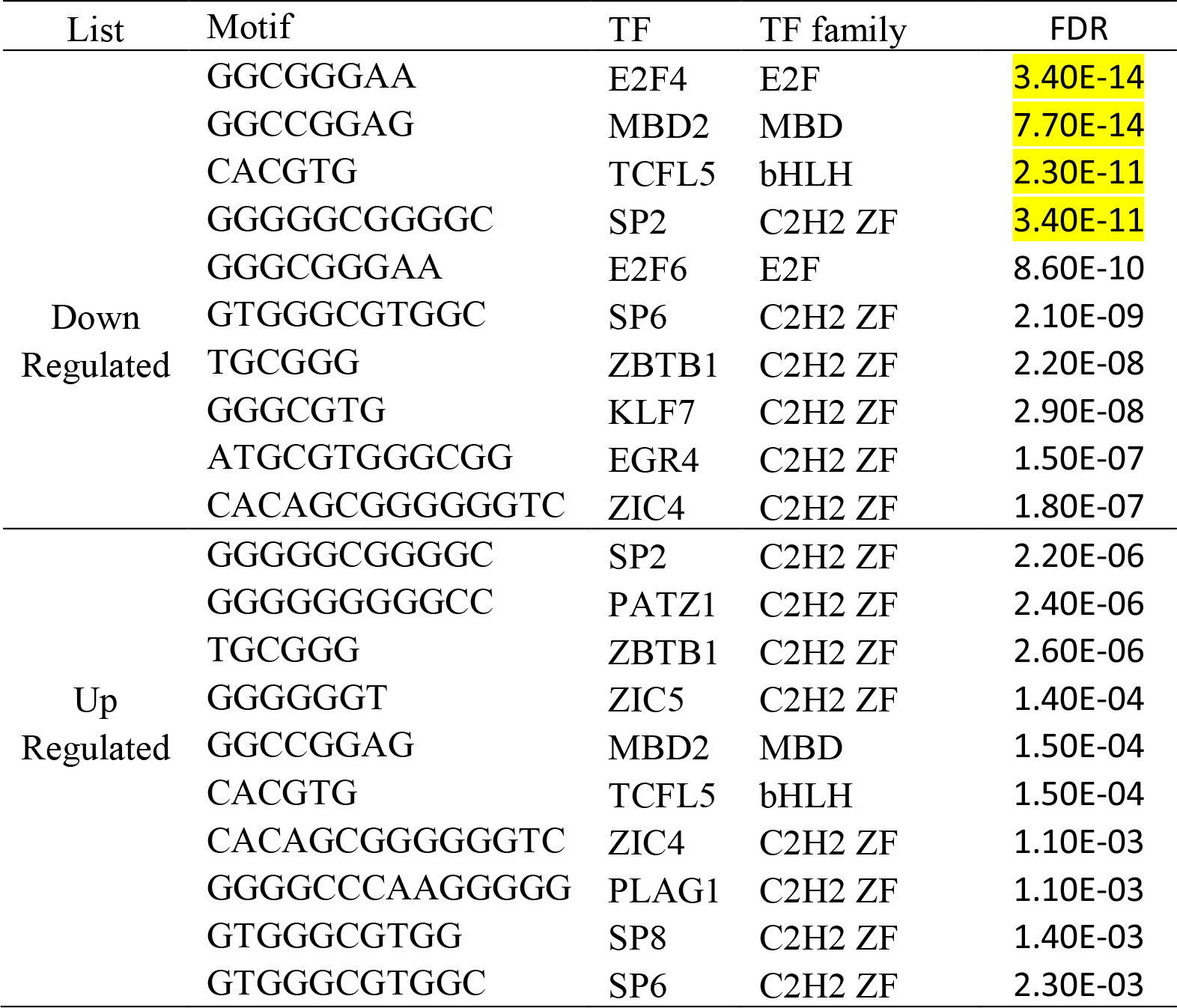
TF motifs enriched in gene promoters (300bp) of up- or down-regulated genes.

For human (Table 2), mouse[25] and Arabidopsis [26], we also include predicted target genes for many miRNAs from multiple sources. Using these gene sets, we detected significant enrichment (Table 6) of miRNA-193b, miR-192, and miR-215 target genes among the down-regulated genes. miR-193b was shown to suppress cell proliferation and down-regulate CCND1 [65]. Proposed as biomarkers of several cancers, miR-192 also inhibit proliferation and can cause cell cycle arrest when overexpressed [66]. miR-215 shares many target genes with miR-192 and are also downregulated in cancer tissues [67]. These miRNAs may play a role in the regulation of cell cycle upon Hoxa1 knockdown.

**Table 6.** Enriched miRNA target gene sets.

| Direction | adj.Pval | nGenes | Pathways |
| --- | --- | --- | --- |
| Down<br>Regulated | 3.40E-45 | 162 | MiRTarBase:miRNA Target hsa-miR-193b-3p |
|  | 1.50E-41 | 170 | MiRTarBase:miRNA Target hsa-miR-192-5p |
|  | 4.20E-41 | 145 | MiRTarBase:miRNA Target hsa-miR-215-5p |
|  | 6.20E-18 | 162 | MiRTarBase:miRNA Target hsa-miR-124-3p |
|  | 3.30E-08 | 82 | MiRTarBase:miRNA Target hsa-miR-34a-5p |
|  | 9.40E-07 | 66 | MiRTarBase:miRNA Target hsa-miR-7-5p |
|  | 9.40E-07 | 58 | MiRTarBase:miRNA Target hsa-miR-375 |
|  | 3.00E-06 | 85 | MiRTarBase:miRNA Target hsa-miR-24-3p |
|  | 4.60E-06 | 89 | MiRTarBase:miRNA Target hsa-miR-1-3p |
|  | 6.80E-05 | 84 | MiRTarBase:miRNA Target hsa-miR-155-5p |
| Up<br>Regulated | 6.00E-16 | 206 | MiRTarBase:miRNA Target hsa-miR-335-5p |
|  | 2.30E-12 | 99 | RegNetwork:miRNA Target hsa-miR-144 |
|  | 1.50E-10 | 81 | RegNetwork:miRNA Target hsa-miR-29c |
|  | 3.30E-10 | 81 | RegNetwork:miRNA Target hsa-miR-29b |
|  | 3.30E-10 | 100 | RegNetwork:miRNA Target hsa-miR-93 |
|  | 3.90E-09 | 72 | RegNetwork:miRNA Target hsa-miR-29a |
|  | 6.20E-09 | 94 | RegNetwork:miRNA Target hsa-miR-30e |
|  | 6.20E-09 | 101 | RegNetwork:miRNA Target hsa-miR-340 |
|  | 6.20E-09 | 67 | RegNetwork:miRNA Target hsa-miR-519d |
|  | 5.70E-08 | 46 | RegNetwork:miRNA Target hsa-miR-17-5p |

#### Pathway analysis

Instead of using selected DEGs that are sensitive to arbitrary cutoffs, pathway analysis can use fold-change values of all genes to identify coherently altered pathways. We used the GAGE [37] as method and KEGG as gene sets. The results (Supp. Table S6) is similar to those from a previous analysis by Turner in an online tutorial [46] and also agrees with our enrichment analysis based on DEGs. For each of the significant KEGG pathways, we can view the fold-changes of related genes on a pathway diagram using the Pathview Bioconductor package [39]. Many cell cycle genes are marked as green in Figure 8, indicating reduced expression in Hoxa1-knockdown samples. We also detected upregulation of genes related to CCRI, arthritis, and lysosome. Many CCR related genes are up-regulated (Figure 9). Not detected using DEGs, lysosome-related genes are mostly upregulated (Supp. Fig. 9). Injected siRNAs might be degraded in the lysosome.

**Figure 9.**
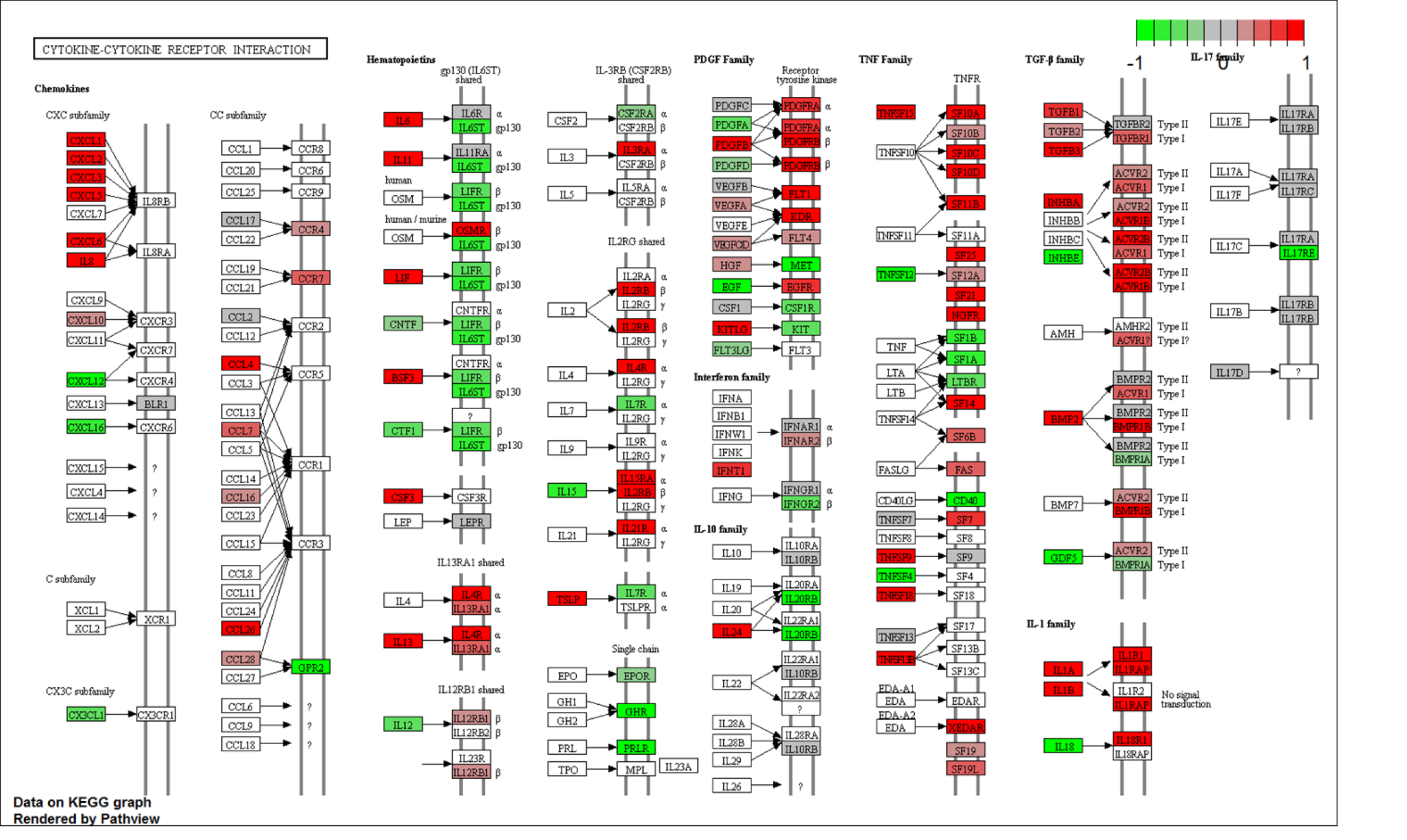
Genes involved in Cytokine-cytokine receptor interaction are mostly up-regulated (red) when Hoxa1 is knocked down. Fold-change information is color-coded on KEGG pathway diagram.

By changing the gene sets database for pathway analysis, we can gain further insights. Using MSigDB.Motif gene sets, we can verify the enrichment of E2F binding motifs (Supp. Table 7). For non-KEGG gene sets, heatmaps are created to show the expression of genes in significant gene sets. Figure 10A shows part of such a plot, highlighting genes that share the “SGCGSSAAA” motif bound by E2F1. Note that E2F1 gene itself is included in the figure, as it binds to its own promoter and forms a positive feedback loop [54]. The downloaded expression data indicate that E2F1 is downregulated by more than 3-fold in Hoxa1 knockdown samples (see Figure 3B). Upon Hoxa1 knockdown, downregulation of E2F1 and downstream genes, including microRNAs, may be part of the transcription program that blocks G_1_/S transition.

**Figure 10.**
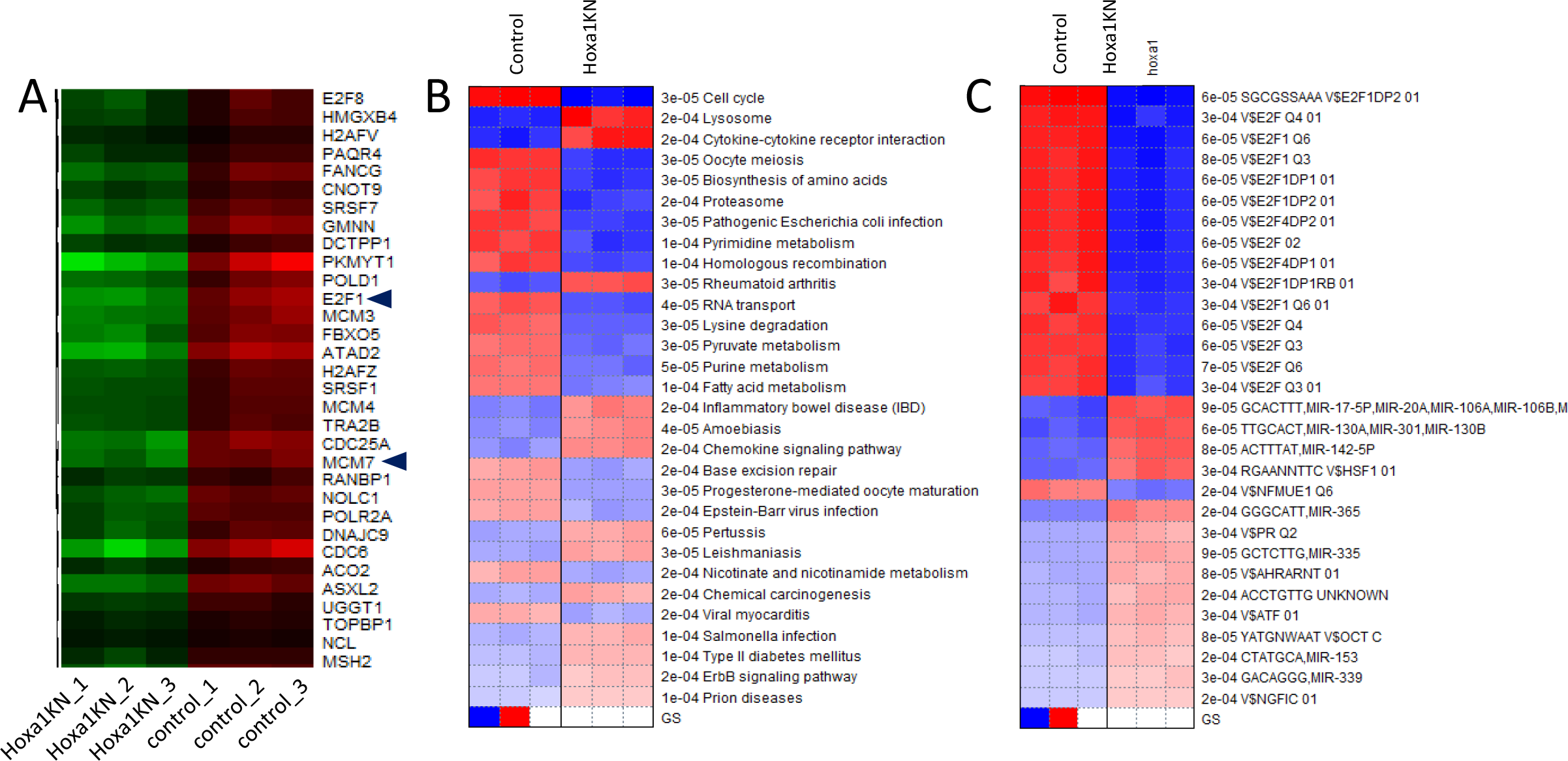
Pathway analysis results using different options. (A) expression patterns of genes with E2F1 binding motifs. E2F1 gene its downregulated in Hoxa1 knockdown. So is the Mcm7 gene, whose intron host miR-106b-25 clusters. (B) Results from running PGSEA sets. (C) PGSEA applied on MSigDB.Motif gene sets.

Users can use many combinations of methods and gene sets to conduct pathway analysis. For example, using PGSEA on KEGG pathways yielded Figure 10B, again confirming previous results on suppressed cell cycle genes and induced lysosome and CCRI related genes. Using the MSigDB.Motif gene sets, we can also confirm the E2F1 binding motifs (Figure 10C). The most highly activated gene sets are related to miR-17-5p, miR-20a, miR106a,b and so on (Figure 10C), which agrees with enrichment analysis using just gene lists.

Some pathways can be attenuated by upregulating some of its associated genes while downregulating others. To detect such pathways, we can use the absolute values of fold changes in pathway analysis. This is achieved by checking the box labeled “Use absolute values of fold changes for GSEA and GAGE.” Instead of detecting up or down-regulated pathways, the results show which pathways are more regulated. As shown in Supp. Table S8, while the expression of ribosome related genes is less variable upon Hoxa1 knockdown, genes related to CCRI are highly regulated.

The expression of neighboring genes can be correlated, due to many mechanisms, including super-enhancers [68], 3D chromatin structure [69], or genomic gain or loss in cancer. To help users detect such correlation, we use ggplot2 [70] and Plotly to interactively visualize fold-changes on all the chromosomes (Figure 11A). Based on regression analysis, the PREDA package [41] can detect statistically significant chromosomal regions with coherent expression change among neighboring genes. Figure 11B shows many such regions in response to Hoxa1 knockdown. Detailed information obtained from downloaded files (Supp. Table S9) suggests, for example, a 4.3 Mbps region on Chr.1q31 contains 6 upregulated genes (PRG4, TPR, C1orf27, PTGS2, PLA2G4A, and BRINP3).

**Figure 11.**
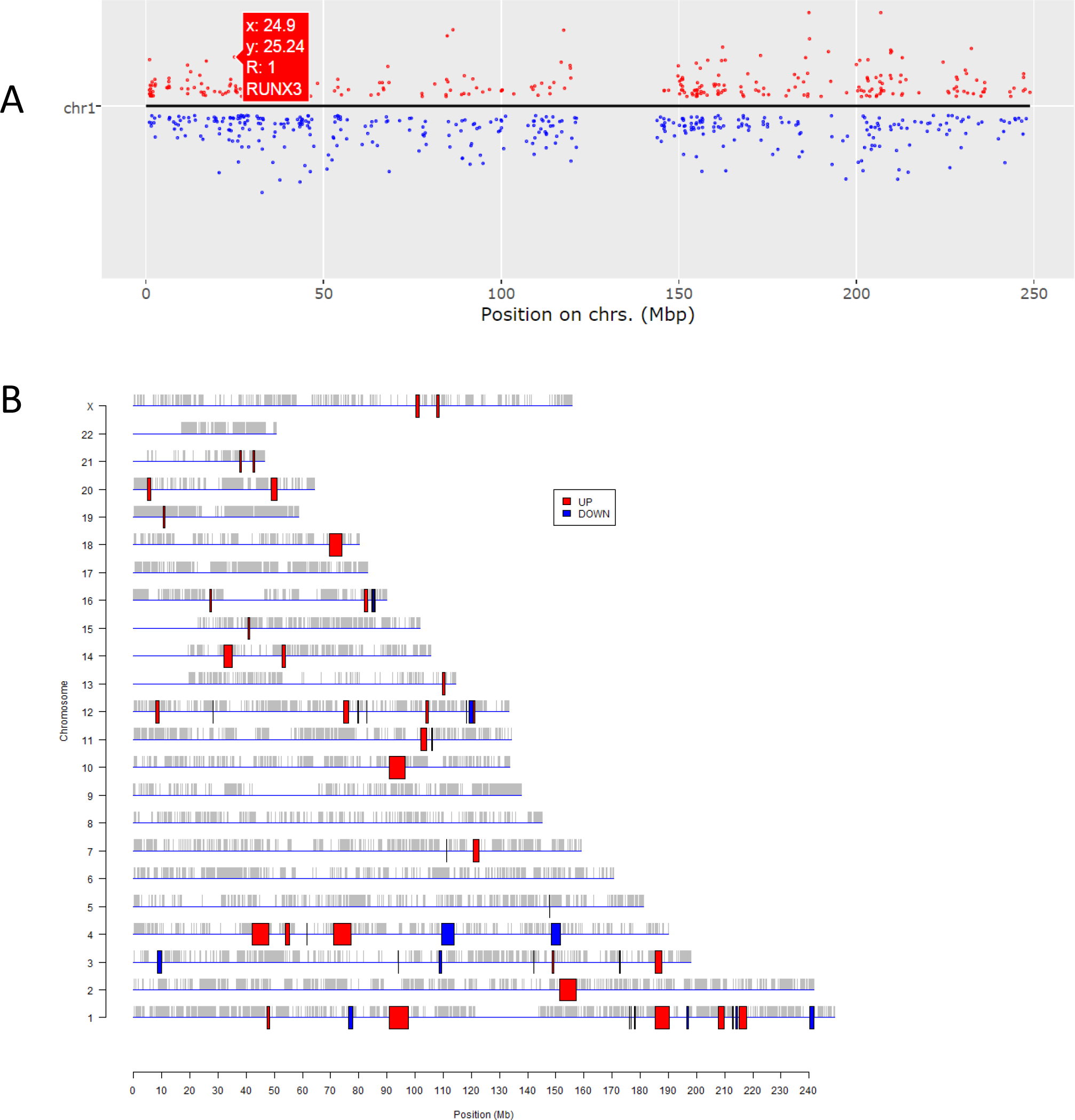
Visualizing expression profiles on chromosomes. A) Zoom-in on Chr. 1 using the dynamic graphics, showing the upregulation of RUX3 gene. B) Statistically significant genomic regions identified by PREDA.

To further validate our parameterization of PREDA, we analyzed DNA microarray data (Supp. File 4) of thymus tissues from patients with Down syndrome [71]. We detected large, upregulated regions on chromosome 21 (Supp. Fig. 10), as expected. Even though PREDA analysis is slow and has low-resolution due to the use of gene-level expression score, it might be useful in cancer studies where localized expression change on the chromosome can happen.

To improve reproducibility, iDEP generates custom R code and R Markdown code based on user data and choices of parameters (Supp. Files 5-7). Users with some R coding experience should be able to re-run most analyses by downloading the related annotation and gene sets used by iDEP. An example is shown here [72].

### Use case 2: factorial design on p53’s role in genotoxic stress

Tonelli *et al*. [29] used RNA-Seq to study the effect of whole-body ionizing radiation (IR) on the mouse with or without p53. B cells and non-B cells were isolated from mouse spleen after treatment. We analyzed the B cell data involving two genotypes (p53 wildtype and p53 null) with mock or IR treatment, a typical 2x2 factorial design. The read count and experimental design files are available as Supp. Files 8 and 9. A converted, filtered version of this dataset is incorporated into iDEP as a demo data.

With this dataset, we demonstrate how users can easily generate hypothesis on molecular pathways and gene regulatory mechanisms through interaction with their data at three main steps: enrichment analysis of k-means clusters, enrichment analysis of the lists of DEGs, and pathway analysis using fold-changes values of all genes.

#### Pre-process and EDA of p53 dataset

We noticed reduced total reads for wildtype samples treated with IR (Figure 12A). While this may be caused by biology, but biased sequencing depth presents a confounding factor, that has not been discussed widely. To quantify such biases, iDEP routinely performs ANOVA analysis of total read counts across sample groups. For this example, uneven read counts are detected (P=0.047) and a warning is produced.

**Figure 12.**
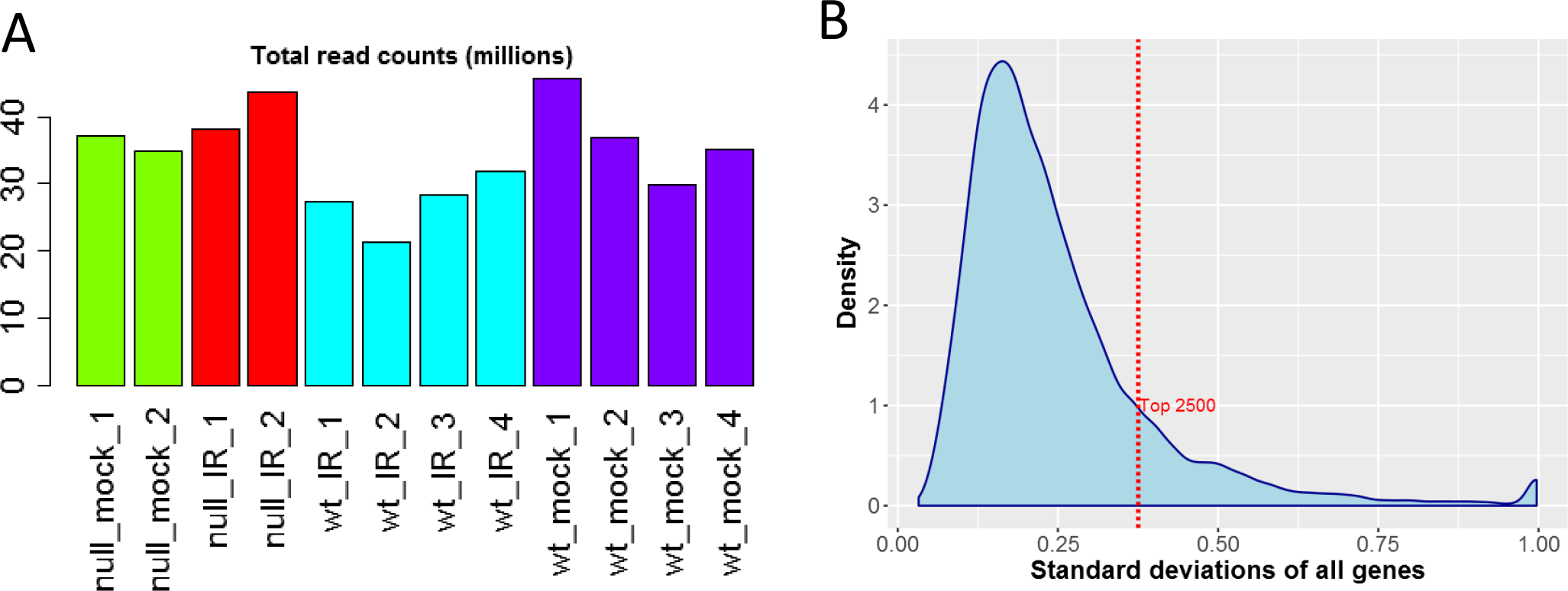
A) Total read counts are smaller in the WT samples treated with IR. B) Distribution of standard deviations for all genes.

EDA shows that IR treatment led to the changes in thousands of genes. Based on the distribution of variances (Figure 12B), we choose the top 2500 genes for clustering analysis. Hierarchical clustering (Figure 13) shows the substantial differences between treated and untreated samples. It also shows the patterns of different groups of genes and the variations among some replicates of treated wild-type cells (wt_IR).

**Figure 13.**
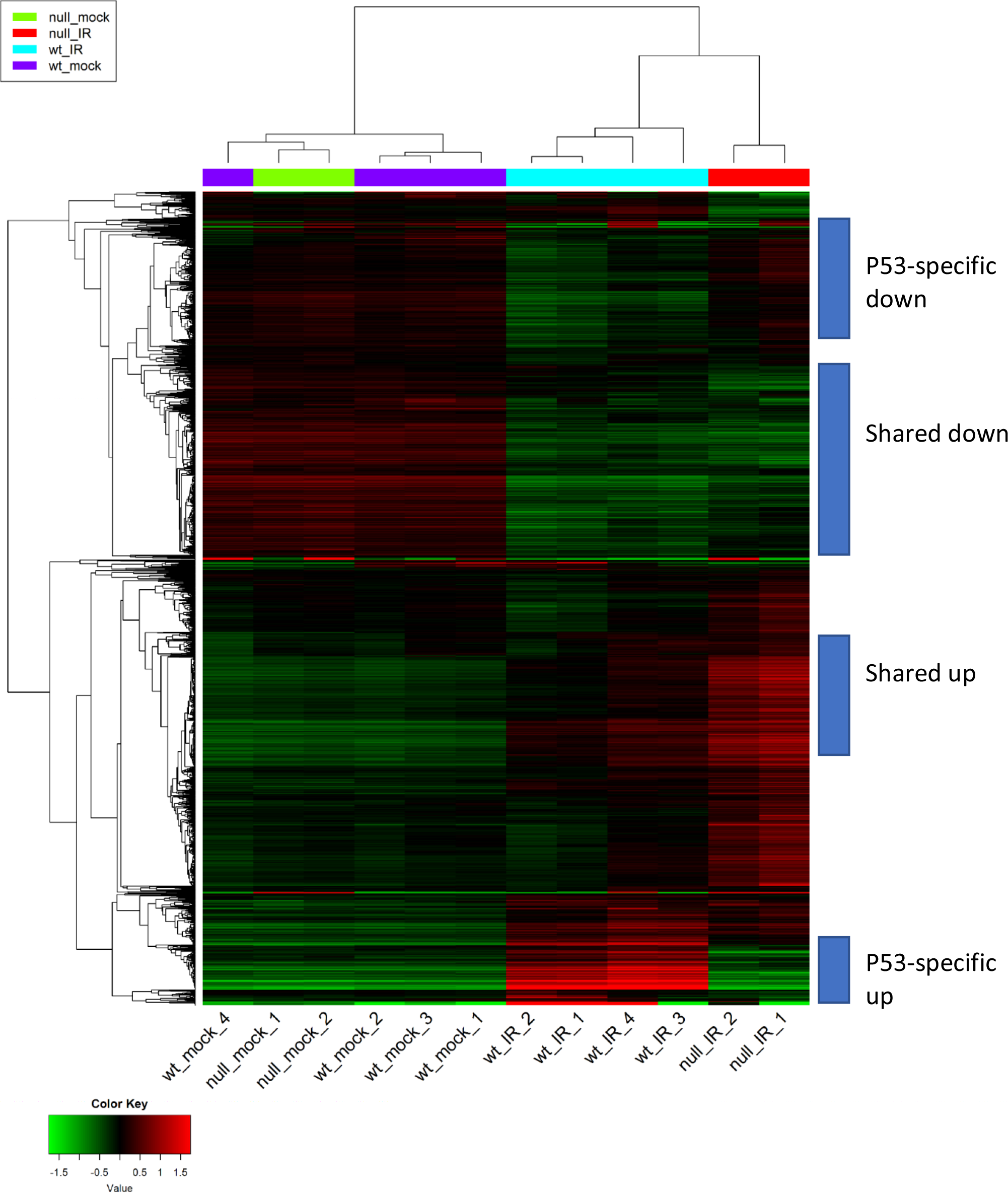
Hierarchical clustering of the 2500 genes shows the patterns of various groups of genes.

We then used k-means clustering to divide the top 2500 genes into groups. Based on the within-group sum of squares plot (Supp. Figure 11) as a reference, we chose a slightly larger k=9. Figure 14 shows the gene clusters and the enriched GO terms. Details are available in Supp. Tables S10 and S11. Genes in clusters B and I show similar responses to IR across genotypes. Strongly enriched in genes related to the immune system (FDR < 3.65×10^−18^), cluster B are downregulated by IR in both cell types. The immune-suppressive effects of radiation [73] are clearly p53-independent. Induced by IR in both wildtype and Trp53^−/−^ cells, cluster I genes are enriched in ribosome biogenesis but with much lower level of significance (FDR < 2.25×10^−5^).

**Figure 14.**
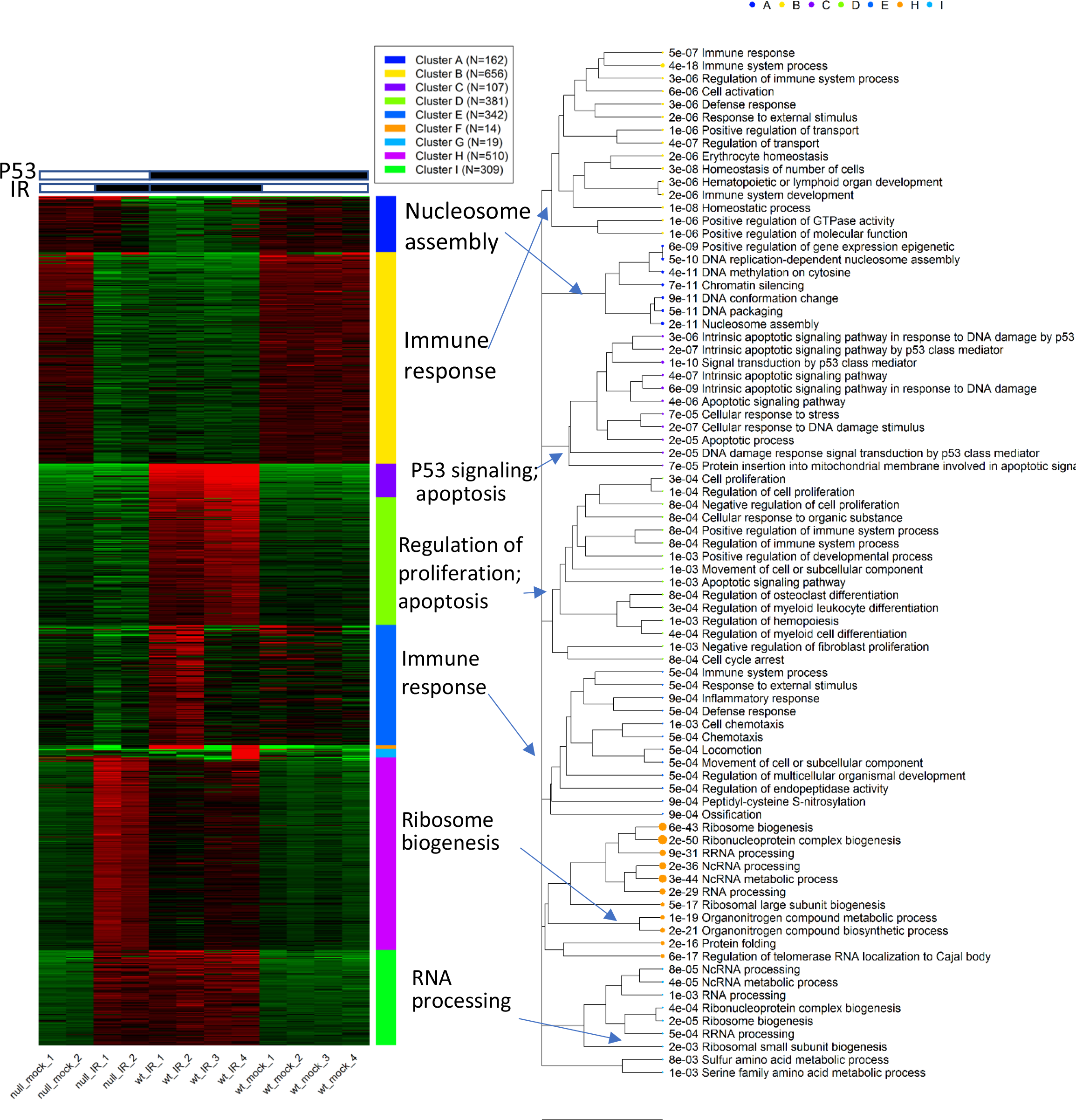
K-means clustering and enrichment analysis.

On the other hand, genes in clusters A, C, and D are specific to the wild-type cells. Cluster A contains 13 genes that code for histone proteins and are involved in nucleosome assembly (FDR<1.66×10^−11^). Genes in Clusters C and D are induced by IR only in B cells with p53, but the former is more strongly upregulated. As expected, cluster C is related to the p53 pathway (FDR<1.38×10^−10^) and apoptosis (FDR<3.59×10^−6^). It is enriched with 15 p53-target genes like Mdm2 (FDR<3.53×10^−18^). Cluster D genes are related to the regulation of cell proliferation and cell cycle arrest, representing further downstream of the transcriptional cascade of p53 signaling.

Genes in cluster H are more highly upregulated in Trp53^−/−^B cells than wildtype cells. It is overrepresented with ncRNA processing (FDR < 3.25×10^−36^), ribosome biogenesis (FDR < 5.53×10^−43^), and protein folding (FDR < 2.23×10^−16^). Many of these genes code for proteins in the nucleus and mitochondrion. Significant enrichment of 7 c-Myc target genes is observed (FDR < 5.09×10^−7^). Many of these enrichment results will be further validated in enrichment analysis of DEGs and pathway analysis. Enrichment analysis of the k-Means clusters provides the first opportunity to gain insight into the molecular pathways underlying different patterns of gene expression.

#### Identifying DEGs in the p53 dataset

To identify genes induced by IR in both cell types, users can use pair-wise comparisons among the 4 sample groups. Alternatively, we can construct linear models through the GUI. Here we use the following model:

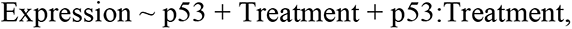

where the last term represents the interaction between genotype and treatment, capturing the additional effects of p53 in IR response. It is important to set the reference levels for factors in a model. Here we set the null (Trp53^−/−^) as a reference level for the factor “p53” and mock for the factor “Treatment”.

With FDR <0.01 and fold-change > 2 as cutoffs, we used DESeq2 to identify DEGs (Figure 15A and B). Without treatment, the two cell types have similar transcription profiles, with few DEGs. But even in Trp53^−/−^ cells, IR caused the upregulation of 1570 genes, 469 of which is also upregulated in p53 wildtype B cells (see Venn diagram in Figure 15C).

**Figure 15.**
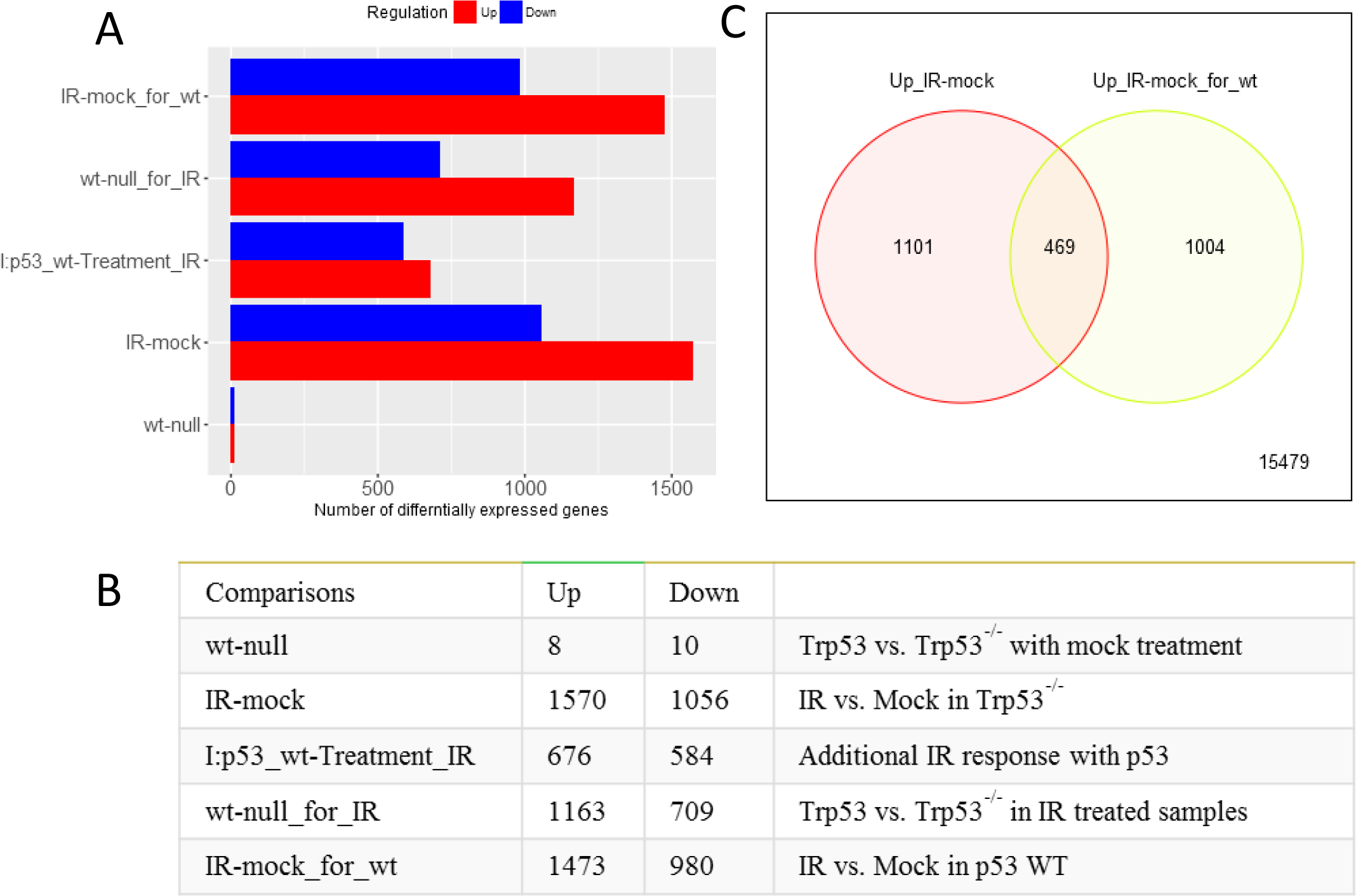
Statistics of DEGs identified by DESeq2. A) and B) shows the numbers of differentially expressed genes for each comparison. C) Venn Diagram shows the overlap between IR induced genes in WT and P53 null samples.

To further understand the molecular pathways, we perform enrichment analysis of the 10 gene lists (Supp. Table S12) associated with 5 comparisons. We focus on two comparisons (1) “IR-mock” representing the baseline response of IR in mutant cells without p53, and (2) “I:p53_wt-Treatment_IR”, the interaction term capturing the additional effect of p53 compared to the baseline response.

For the first comparison, Figure 16 shows IR induced DEGs in mutant cells. The 1570 upregulated genes are related to non-coding RNA (ncRNA) metabolic process (FDR <1.33×10^−79^), ribosome biogenesis (FDR<2.54×10^−67^), and translation (FDR <3.03×10^−32^). This enrichment profile is similar to cluster H derived from the k-Means clustering, as the two lists capture the same group of genes. The upregulated genes are surprisingly coherent in function. For example, 219 (14%) can be found in the nucleus, 286 (18%) is related to the mitochondrion, and, most significantly, 407 (26%) is RNA-binding (FDR<3.54×10^−138^). The 1570 upregulated genes contain 7 MYC target genes (FDR<4.22×10^−7^), consistent with the fact that MYC is a direct regulator of ribosome biogenesis[74]. This agrees with reports of the involvement of MYC in radiation treatment [75, 76], suggesting MYC may trigger proliferation pathways upon genotoxic stress, in the absence of p53.

**Figure 16.**
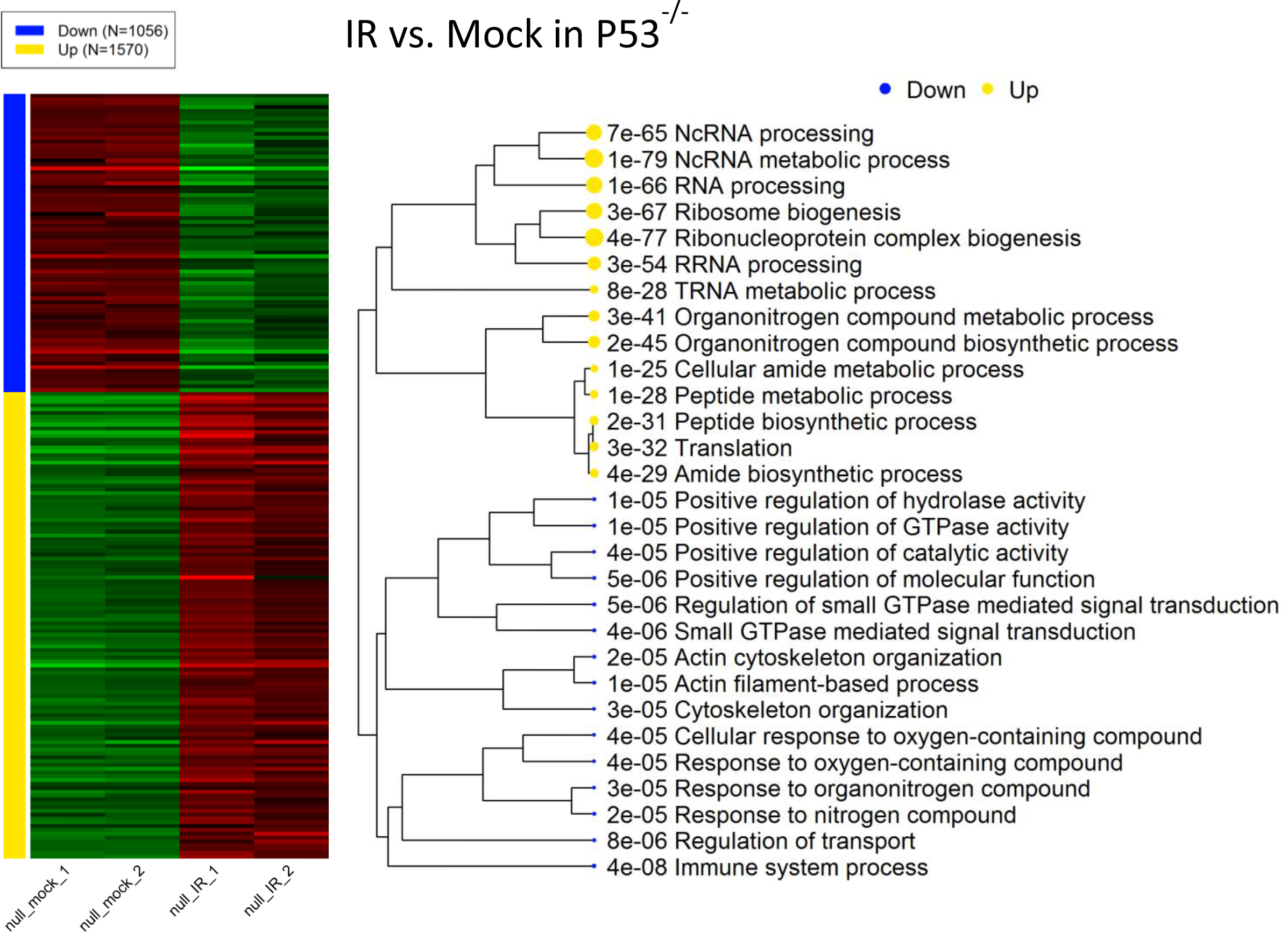
Effect of IR on p53 null samples. A) The expression patterns of DEGs. B) Up-regulated genes are highly overrepresented with Rib genesis, RNA, especially ncRNA, processing, and translation. Downregulated genes are enriched genes involved with immune system, a oskeleton, and GTPase activity.

Genes downregulated by IR in Trp53^−/−^ B cells are related to immune system (FDR< 4.22×10^−8^), GTPase activity (FDR < 3.75×10^−6^), and actin cytoskeleton (FDR < 2.06×10^−5^). As shown in Supp. Table S13, we can also detect the enrichment of the target genes of miR-124 (FDR< 4.56×10^−12^), an important modulator of immunity [77]. Others associated miRNAs, including miR-6931-5p, Mir-4321, and miR-576-5p, may also be involved.

For the second comparison, the expression profiles of DEGs associated with the interaction term is shown in Figure 17. This is the p53 mediated IR response, compared to the baseline response without p53. The 676 genes that are upregulated in wild-type B cells following IR, but not in Trp53^−/−^ B cells. As expected, these genes are enriched in p53-mediated response to DNA damage (FDR< 1.43×10^−6^), and apoptosis (FDR < 9.72×10^−6^). As shown in Supp. Table S13, these genes are overrepresented with 25 target genes of p53 (FDR < 1.34×10^−13^) and 76 target genes of miR-92a (FDR < 2.79×10^−11^). Part of the miR-17/92 cluster, miR-92a is related to tumorigenesis and is regulated by p53 [78, 79]. Another miRNA with overrepresented target genes is miR-504 (FDR < 3.25×10^−8^), which has been shown to binds to 3’ UTR of Trp53 and negatively regulate its expression [80]. Located in the introns of the fibroblast growth factor 13 (FGF13) gene, miR-504 is transcriptionally suppressed by p53, forming a negative feedback loop [81]. Following radiation, the expression of both miR-92a and miR-504 in wild-type B cells may be reduced, leading to the upregulation of their target genes. Further study is needed to verify this hypothesis.

**Figure 17.**
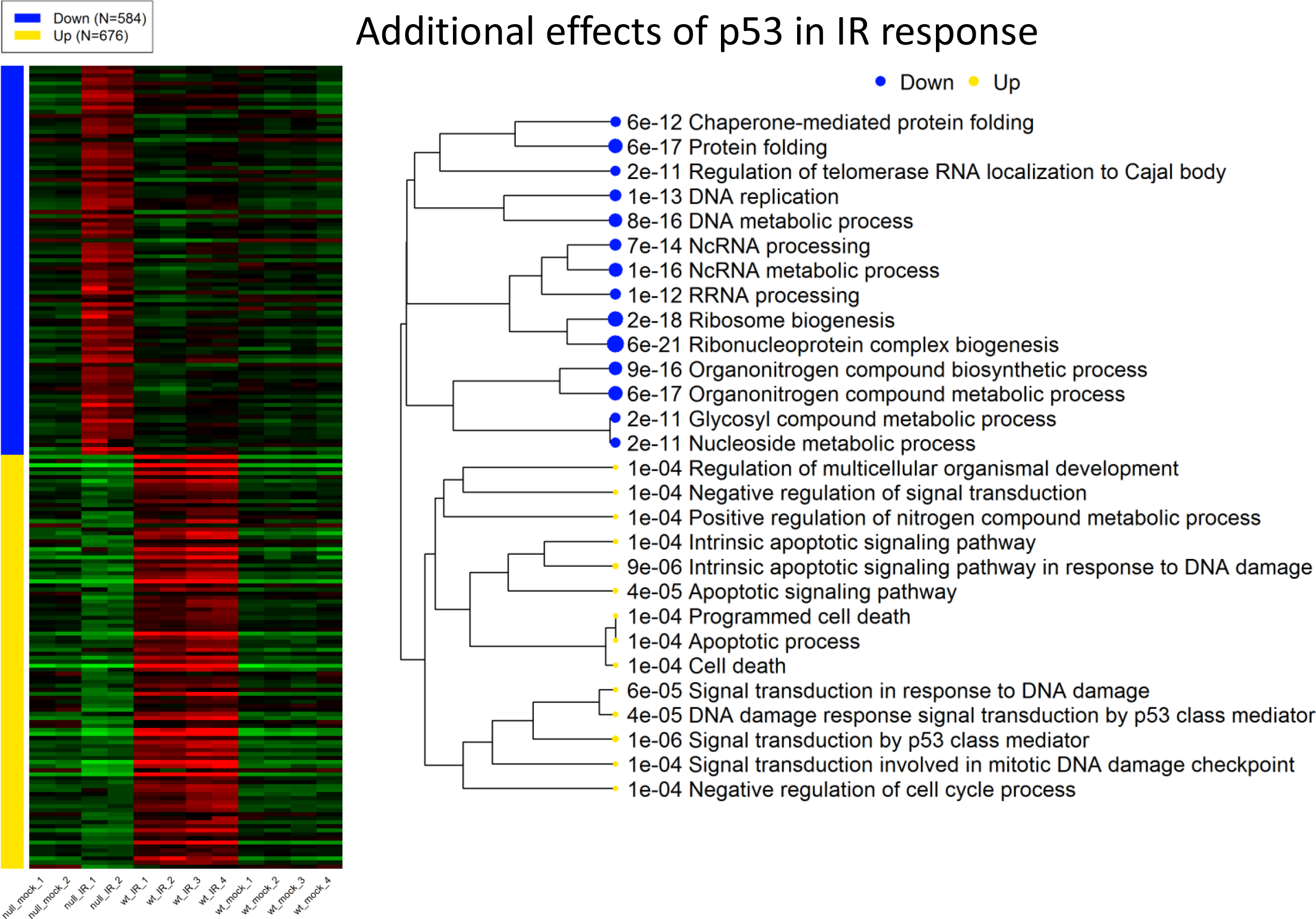
Additional effect of p53 in IR response. A) Expression patterns of selected DEGs. B) Upregulated genes are enriched with genesd to p53 mediated response to DNA damage, especially apoptosis, and negative regulation of cell cycle. These genes are only induced b ells with wildtype p53. One the other hand, p53 caused the relative downregulation of genes related to ribosome biogenesis, tRNA and processing, DNA replication, and protein folding. These genes are only upregulated by IR in Trp53^−/−^.

As shown in Figure 17, the 584 genes downregulated according to the interaction term are those that are induced in the Trp53^−/−^ B cells, but not in wild-type B cells. These genes are overrepresented with ncRNA processing, ribosome biogenesis, cell cycle, and RNA transport (Supp. Table S14). Most (411) of the 584 genes are included in the genes upregulated by IR in Trp53^−/−^ B cells (Supp. Figure 12). MYC target genes are also downregulated by p53 upon IR. In wildtype B cells, p53 suppresses the MYC oncogenic pathway compared to Trp53^−/−^ B cells. The most significant shared TF binding motif is E2F1 (FDR < 7.73×10^−11^). This agrees with the role of p53 in cell cycle arrest through p21-mediated control of E2F factors [82].

#### Pathway analysis of p53 data

Many of the above observations can be confirmed by using pathway analysis (Supp. Table S15) based on the fold-change values of all genes. The results of GSEA on the interaction term can be found in Supp. Table S15. The PGSEA package offers a convenient way to visualize the activities of pathways across all samples. Figure 18 clearly shows that p53 signaling pathway, apoptosis, and positive regulation of cell cycle arrest are uniquely activated by IR in wild-type B cells. This is again confirmed by TF target genes (Figure 19). In addition, the p53-independent upregulation of MYC target genes can also be observed in Figure 19. Several ETS transcription factors, including SFPI1, SPI1, and ETS1, are suppressed by IR in both cell types. These factors may underlie the suppression of immune response as suggested [83]. Applying PGSEA on miRNA target genes highlights miRNA-30a (Supp. Figure 15), whose target genes are specifically activated by IR in wild-type B cells. miRNA-30a was shown to be involved in response to IR [84] and mutually regulate p53 [85]. Thus, the complex p53 signaling pathways are unveiled with remarkable accuracy.

**Figure 18.**
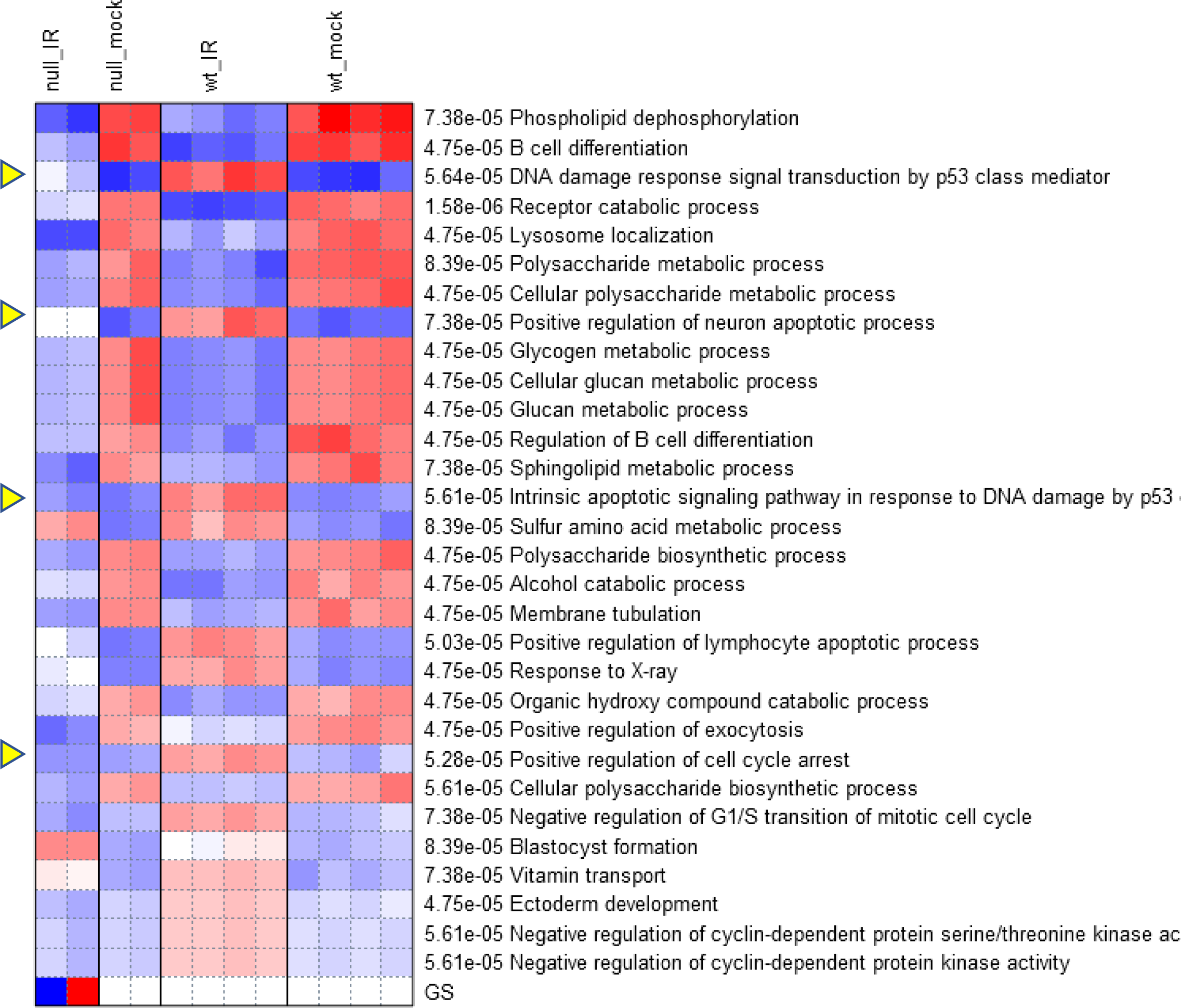
PGSEA visualization of pathway activities using GO biological processes.

**Figure 19.**
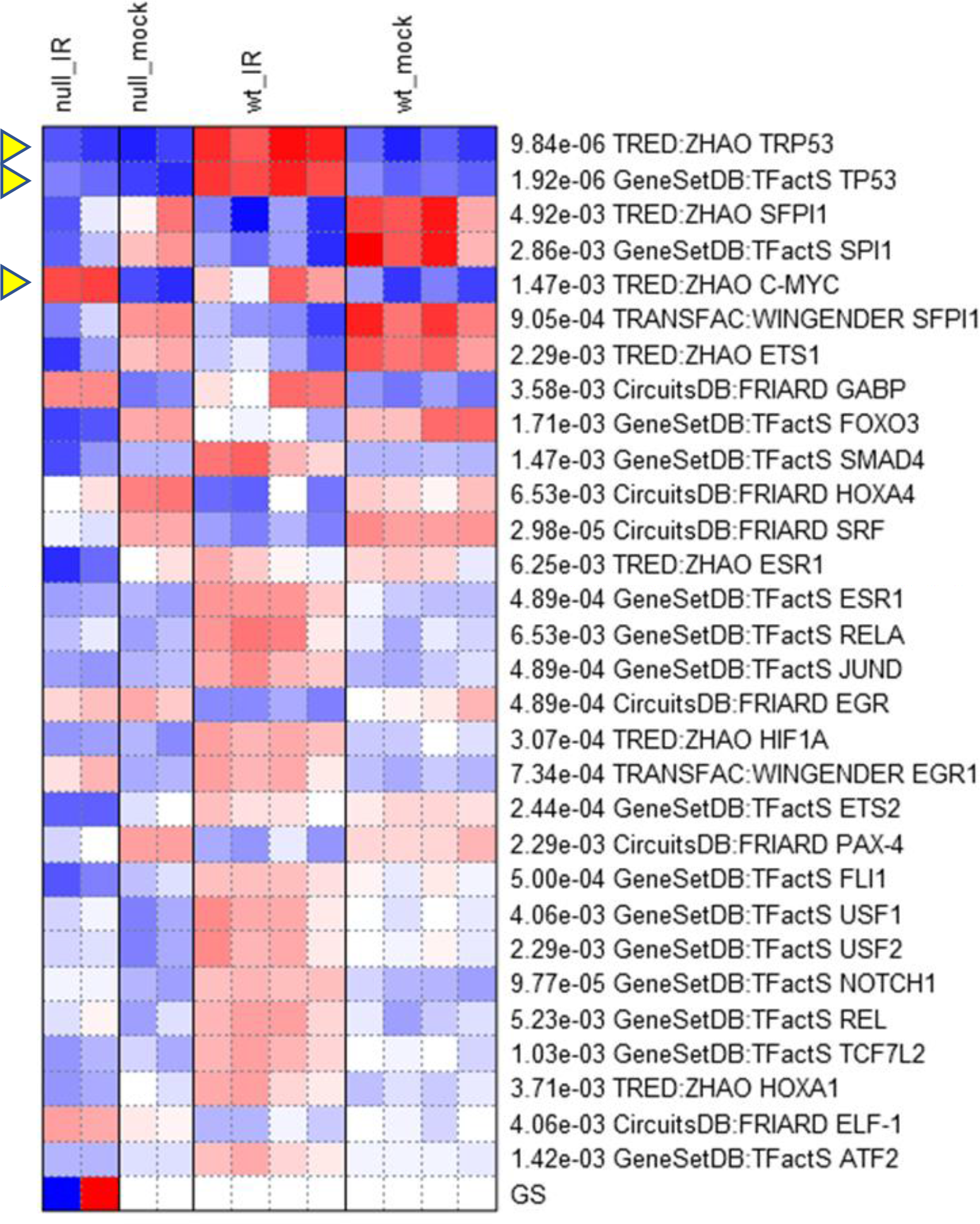
Differentially regulated TF target gene sets across sample types, identified by PGSEA.

The upregulated p53 target genes can be seen in the KEGG pathway diagram (Figure 20). This pathway map shows multifaceted roles of p53 in the regulation of apoptosis, cell cycle, DNA damage repair, and growth arrest. Many of these functions were re-discovered in our analyses above. This shows the power of comprehensive pathway databases coupled with broad analytic functionalities accessible via an intuitive user interface. Without iDEP, it can take days or weeks to write code and collect data to conduct all the analyses above. With iDEP, biologists can complete such analyses in as little as 20 minutes.

**Figure 20.**
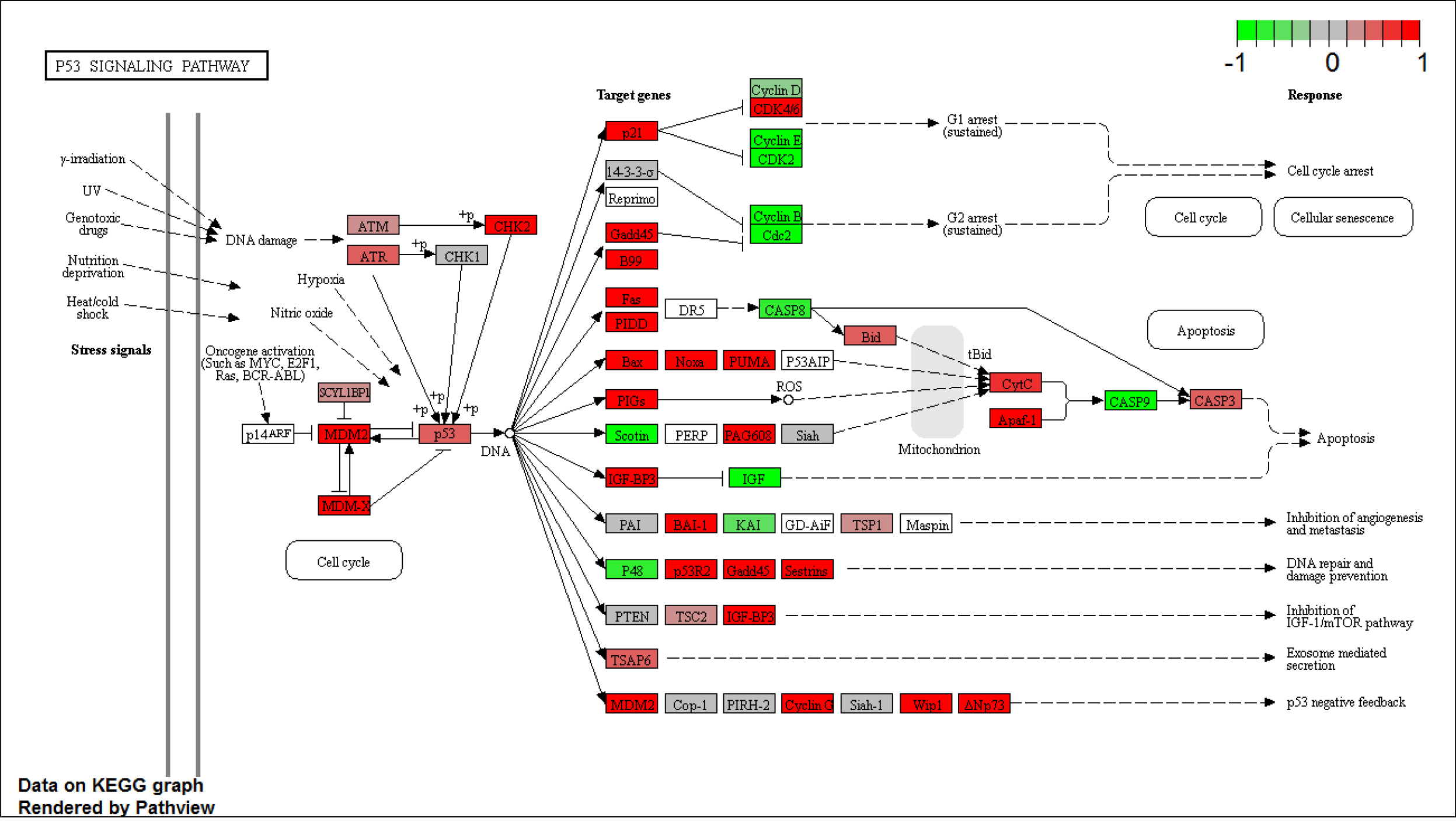
Fold changes of genes in P53 signaling pathway in wildtype B cells treated with IR. Upregulation (red) of many key-players.

## Discussions and conclusions

By integrating many Bioconductor packages with comprehensive annotation databases, iDEP enables users to conduct in-depth bioinformatics analysis of transcriptomic data through a GUI. Taking advantage of the Shiny platform, we were able to pack many useful functionalities into iDEP, including high-quality graphics based on ggplot2 and interactive plots using Plotly. Compared with traditional web applications, Shiny has its drawbacks and limitations. The interface is not as flexible as those developed using JavaScript.

Nevertheless, we believe an integrated web application like iDEP is a valuable tool to both bench scientists and bioinformaticians.

As an example, we extensively analyzed an RNA-Seq dataset involving Hoxa1 knockdown by siRNA in lung fibroblasts, and identified the down-regulation of cell-cycle genes, in agreement with previous analyses and experimental confirmation. Our analyses also show E2F and SP1 binding motifs are enriched in the promoters of downregulated genes, mediating the cell cycle arrest. Furthermore, we also find evidence that microRNAs (miR-17-5P, miR-20a, miR-106a, miR-192, miRNA-193b, and miR-215) might work together with E2F factors to block the G_1_/S transition in response to reduced Hoxa1 expression. Interestingly, miR-106a is located in the intron of Mcm7, an E2F1 target gene. DEGs are also enriched with genes related to neuron parts, synapse, as well as neurodegenerative diseases. This is consistent with reports of Hoxa1’s role in neuron differentiation [48-50]. Hoxa1 knockdown induces expression of genes associated with the cytokine-cytokine interaction, lysosome, and cell migration, probably in response to the injected siRNAs. These genes are overrepresented with target genes of NF-κB, known to be involved in immune response. By combining both annotation dataset and analytic functionality, iDEP help biologists to quickly analyze their data from different perspectives.

In the second example, our analysis shows that in B cell without p53, radiation treatment upregulates MYC oncogenic pathway, triggering downstream genes with highly coherent functions such as cell proliferation, ribosome biogenesis, and ncRNA metabolism. Enriched with target genes of miR-124, and ETS domain transcription factors, genes downregulated by IR in p53 null B cells are associated with immune response, GTPase activity and actin cytoskeleton. In wildtype B cells, a p53-dependent transcriptional response to IR is evidently related to p53-mediated apoptosis and DNA repair, as expected. The target genes of MYC and E2F1 are suppressed by p53, leading to growth and cell cycle arrest. iDEP helps unveil the multifaceted functions of p53, and also highlight the potential involvement of several miRNAs (miR-92a, miR-504, and miR-30a).

Users should be cautious when interpreting results from pathway analysis, which can be obtained through the many combinations of methods and gene set databases. The biomedical literature is large and heterogeneous [86], making it easy to rationalize and make a story out of any gene. True pathways, like the effect of Hoxa1 knockdown on cell cycle, should be robustly identified across different methods and databases. Also, as demonstrated in the two examples, for each enrichment or pathway analysis, we tried to focus on the most significant gene sets.

Besides RNA-Seq and DNA microarray data, users can also use iDEP to analyze fold-change and FDR values calculated by other methods such as cuffdiff [87]. For unannotated genomes, iDEP can only be used for EDA and differential expression analysis. For single-cell RNA-Seq data [88], only smaller, pre-processed, imputed datasets with hundreds of cells can be analyzed, as iDEP is mostly designed to handle transcriptomic data derived from bulk tissues.

In addition to updating the annotation database derived from Ensembl every year, we plan to continue to compile pathway databases for model organisms, similar to MSigDB and GSKB. For currently unsupported species, we will consider ways to incorporate user-submitted gene annotation. Based on user request and feedback, we will also add more functions by including additional Bioconductor packages.

## Acknowledgments

The author thanks Brian Moore and Kevin Brandt for technical support of the web server. iDEP is strengthened by constructive criticisms from a previous reviewer, and many suggestions from users.

## References

1. Mortazavi A, Williams BA, McCue K, Schaeffer L, Wold B: Mapping and quantifying mammalian transcriptomes by RNA-Seq. Nat Methods 2008, 5:621–628.

2. Kim D, Pertea G, Trapnell C, Pimentel H, Kelley R, Salzberg SL: TopHat2: accurate alignment of transcriptomes in the presence of insertions, deletions and gene fusions. Genome Biol 2013, 14:R36.

3. Trapnell C, Hendrickson DG, Sauvageau M, Goff L, Rinn JL, Pachter L: Differential analysis of gene regulation at transcript resolution with RNA-seq. Nat Biotechnol 2013, 31:46–53.

4. Bray NL, Pimentel H, Melsted P, Pachter L: Near-optimal probabilistic RNA-seq quantification. Nat Biotechnol 2016, 34:525–527.

5. Patro R, Duggal G, Love MI, Irizarry RA, Kingsford C: Salmon provides fast and bias-aware quantification of transcript expression. Nat Methods 2017, 14:417–419.

6. Reich M, Liefeld T, Gould J, Lerner J, Tamayo P, Mesirov JP: GenePattern 2.0. Nat Genet 2006, 38:500–501.

7. Afgan E, Baker D, van den Beek M, Blankenberg D, Bouvier D, Cech M, Chilton J, Clements D, Coraor N, Eberhard C, et al: The Galaxy platform for accessible, reproducible and collaborative biomedical analyses: 2016 update. Nucleic Acids Res 2016, 44:W3–W10.

8. Merchant N, Lyons E, Goff S, Vaughn M, Ware D, Micklos D, Antin P: The iPlant Collaborative: Cyberinfrastructure for Enabling Data to Discovery for the Life Sciences. PLoS Biol 2016, 14:e1002342.

9. Huber W, Carey VJ, Gentleman R, Anders S, Carlson M, Carvalho BS, Bravo HC, Davis S, Gatto L, Girke T, et al: Orchestrating high-throughput genomic analysis with Bioconductor. Nat Methods 2015, 12:115–121.

10. Love MI, Huber W, Anders S: Moderated estimation of fold change and dispersion for RNA-seq data with DESeq2. Genome Biol 2014, 15:550.

11. Nelson JW, Sklenar J, Barnes AP, Minnier J: The START App: a web-based RNAseq analysis and visualization resource. Bioinformatics 2016.

12. Dai Z, Sheridan JM, Gearing LJ, Moore DL, Su S, Wormald S, Wilcox S, O’Connor L, Dickins RA, Blewitt ME, Ritchie ME: edgeR: a versatile tool for the analysis of shRNA-seq and CRISPR-Cas9 genetic screens. F1000Res 2014, 3:95.

13. Law CW, Chen Y, Shi W, Smyth GK: voom: Precision weights unlock linear model analysis tools for RNA-seq read counts. Genome Biol 2014, 15:R29.

14. Pimentel H, Bray N, Puente S, Melsted P, Pachter L: Differential analysis of RNA-Seq incorporating quantification uncertainty. In BioRxiv, vol. http://dx.doi.org/10.1101/058164; 2016.

15. Harshbarger J, Kratz A, Carninci P: DEIVA: a web application for interactive visual analysis of differential gene expression profiles. BMC Genomics 2017, 18:47.

16. Younesy H, Moller T, Lorincz MC, Karimi MM, Jones SJ: VisRseq: R-based visual framework for analysis of sequencing data. BMC Bioinformatics 2015, 16 Suppl 11:S2.

17. Gardeux V, David F, Shajkofci A, Schwalie P, Deplancke B: ASAP: A web-based platform for the analysis and interactive visualization of single-cell RNA-seq data In bioRxiv. pp. 096222; 2016:096222.

18. Ashburner M, Ball CA, Blake JA, Botstein D, Butler H, Cherry JM, Davis AP, Dolinski K, Dwight SS, Eppig JT, et al: Gene ontology: tool for the unification of biology. The Gene Ontology Consortium. Nat Genet 2000, 25:25–29.

19. Kanehisa M, Furumichi M, Tanabe M, Sato Y, Morishima K: KEGG: new perspectives on genomes, pathways, diseases and drugs. Nucleic Acids Res 2017, 45:D353–D361.

20. Zheng HQ, Wu NY, Chow CN, Tseng KC, Chien CH, Hung YC, Li GZ, Chang WC: EXPath tool-a system for comprehensively analyzing regulatory pathways and coexpression networks from high-throughput transcriptome data. DNA Res 2017.

21. Szklarczyk D, Franceschini A, Wyder S, Forslund K, Heller D, Huerta-Cepas J, Simonovic M, Roth A, Santos A, Tsafou KP, et al: STRING v10: protein-protein interaction networks, integrated over the tree of life. Nucleic Acids Res 2015, 43:D447–452.

22. Aken BL, Achuthan P, Akanni W, Amode MR, Bernsdorff F, Bhai J, Billis K, Carvalho-Silva D, Cummins C, Clapham P, et al: Ensembl 2017. Nucleic Acids Res 2017, 45:D635–D642.

23. Aken BL, Ayling S, Barrell D, Clarke L, Curwen V, Fairley S, Fernandez Banet J, Billis K, Garcia Giron C, Hourlier T, et al: The Ensembl gene annotation system. Database (Oxford) 2016, 2016.

24. Bolser DM, Staines DM, Perry E, Kersey PJ: Ensembl Plants: Integrating Tools for Visualizing, Mining, and Analyzing Plant Genomic Data. Methods Mol Biol 2017, 1533:1–31.

25. Lai ea: GSKB: A gene set database for pathway analysis in mouse. bioRxiv 2016, 0802511.

26. Lai L, Liberzon A, Hennessey J, Jiang G, Qi J, Mesirov JP, Ge SX: AraPath: a knowledgebase for pathway analysis in Arabidopsis. Bioinformatics 2012, 28:2291–2292.

27. Liberzon A, Birger C, Thorvaldsdottir H, Ghandi M, Mesirov JP, Tamayo P: The Molecular Signatures Database (MSigDB) hallmark gene set collection. Cell Syst 2015, 1:417–425.

28. Fabregat A, Sidiropoulos K, Garapati P, Gillespie M, Hausmann K, Haw R, Jassal B, Jupe S, Korninger F, McKay S, et al: The Reactome pathway Knowledgebase. Nucleic Acids Res 2016, 44:D481–487.

29. Tonelli C, Morelli MJ, Bianchi S, Rotta L, Capra T, Sabo A, Campaner S, Amati B: Genome-wide analysis of p53 transcriptional programs in B cells upon exposure to genotoxic stress in vivo. Oncotarget 2015, 6:24611–24626.

30. van der Maaten LJP, Hinton GE: Visualizing High-Dimensional Data Using t-SNE. Journal of Machine Learning Research 2008, 9:2579–2605.

31. Ritchie ME, Phipson B, Wu D, Hu Y, Law CW, Shi W, Smyth GK: limma powers differential expression analyses for RNA-sequencing and microarray studies. Nucleic Acids Res 2015, 43:e47.

32. Ge SX: Exploratory bioinformatics investigation reveals importance of “junk” DNA in early embryo development. BMC Genomics 2017, 18:200.

33. Liu R, Holik AZ, Su S, Jansz N, Chen K, Leong HS, Blewitt ME, Asselin-Labat ML, Smyth GK, Ritchie ME: Why weight? Modelling sample and observational level variability improves power in RNA-seq analyses. Nucleic Acids Res 2015, 43:e97.

34. Subramanian A, Tamayo P, Mootha VK, Mukherjee S, Ebert BL, Gillette MA, Paulovich A, Pomeroy SL, Golub TR, Lander ES, Mesirov JP: Gene set enrichment analysis: a knowledge-based approach for interpreting genome-wide expression profiles. Proc Natl Acad Sci U S A 2005, 102:15545–15550.

35. Kim SY, Volsky DJ: PAGE: parametric analysis of gene set enrichment. BMC Bioinformatics 2005, 6:144.

36. Furge K, Dykema K: PGSEA: Parametric Gene Set Enrichment Analysis. R package version 1480 2012.

37. Luo W, Friedman MS, Shedden K, Hankenson KD, Woolf PJ: GAGE: generally applicable gene set enrichment for pathway analysis. BMC Bioinformatics 2009, 10:161.

38. Yu G, He QY: ReactomePA: an R/Bioconductor package for reactome pathway analysis and visualization. Mol Biosyst 2016, 12:477–479.

39. Luo W, Brouwer C: Pathview: an R/Bioconductor package for pathway-based data integration and visualization. Bioinformatics 2013, 29:1830–1831.

40. Araki H, Knapp C, Tsai P, Print C: GeneSetDB: A comprehensive meta-database, statistical and visualisation framework for gene set analysis. FEBS Open Bio 2012, 2:76–82.

41. Ferrari F, Solari A, Battaglia C, Bicciato S: PREDA: an R-package to identify regional variations in genomic data. Bioinformatics 2011, 27:2446–2447.

42. Kluger Y, Basri R, Chang JT, Gerstein M: Spectral biclustering of microarray data: coclustering genes and conditions. Genome Res 2003, 13:703–716.

43. Zhang Y, Xie J, Yang J, Fennell A, Zhang C, Ma Q: QUBIC: a bioconductor package for qualitative biclustering analysis of gene co-expression data. Bioinformatics 2017, 33:450–452.

44. Orzechowski P, Panszczyk A, Huang XY, Moore JH: runibic: a Bioconductor package for parallel row-based biclustering of gene expression data. In BioRxiv, vol. 2017. pp. 210682; 2017:210682.

45. Langfelder P, Horvath S: WGCNA: an R package for weighted correlation network analysis. BMC Bioinformatics 2008, 9:559.

46. Turner S: Tutorial: RNA-seq differential expression&pathway analysis with Sailfish, DESeq2, GAGE, and Pathview. http://www.gettinggeneticsdone.com/2015/12/tutorial-rna-seq-differential.html; 2015.

47. Jung D, Ge SX: PPInfer: a Bioconductor package for inferring functionally related proteins using protein interaction networks. F1000Research 2018, 6:1969.

48. Paraguison RC, Higaki K, Yamamoto K, Matsumoto H, Sasaki T, Kato N, Nanba E: Enhanced autophagic cell death in expanded polyhistidine variants of HOXA1 reduces PBX1-coupled transcriptional activity and inhibits neuronal differentiation. J Neurosci Res 2007, 85:479–487.

49. Gavalas A, Ruhrberg C, Livet J, Henderson CE, Krumlauf R: Neuronal defects in the hindbrain of Hoxa1, Hoxb1 and Hoxb2 mutants reflect regulatory interactions among these Hox genes. Development 2003, 130:5663–5679.

50. Canu E, Boccardi M, Ghidoni R, Benussi L, Duchesne S, Testa C, Binetti G, Frisoni GB: HOXA1 A218G polymorphism is associated with smaller cerebellar volume in healthy humans. J Neuroimaging 2009, 19:353–358.

51. Ge SX: Large-scale analysis of expression signatures reveals hidden links among diverse cellular processes. BMC Syst Biol 2011, 5:87.

52. Vermeulen K, Van Bockstaele DR, Berneman ZN: The cell cycle: a review of regulation, deregulation and therapeutic targets in cancer. Cell Prolif 2003, 36:131–149.

53. Nahle Z, Polakoff J, Davuluri RV, McCurrach ME, Jacobson MD, Narita M, Zhang MQ, Lazebnik Y, Bar-Sagi D, Lowe SW: Direct coupling of the cell cycle and cell death machinery by E2F. Nat Cell Biol 2002, 4:859–864.

54. DeGregori J: The genetics of the E2F family of transcription factors: shared functions and unique roles. Biochim Biophys Acta 2002, 1602:131–150.

55. Motokura T, Arnold A: PRAD1/cyclin D1 proto-oncogene: genomic organization, 5’ DNA sequence, and sequence of a tumor-specific rearrangement breakpoint. Genes Chromosomes Cancer 1993, 7:89–95.

56. Grinstein E, Jundt F, Weinert I, Wernet P, Royer HD: Sp1 as G1 cell cycle phase specific transcription factor in epithelial cells. Oncogene 2002, 21:1485–1492.

57. Lin SY, Black AR, Kostic D, Pajovic S, Hoover CN, Azizkhan JC: Cell cycle-regulated association of E2F1 and Sp1 is related to their functional interaction. Mol Cell Biol 1996, 16:1668–1675.

58. Baeuerle PA, Henkel T: Function and activation of NF-kappa B in the immune system. Annu Rev Immunol 1994, 12:141–179.

59. Dejean AS, Beisner DR, Ch’en IL, Kerdiles YM, Babour A, Arden KC, Castrillon DH, DePinho RA, Hedrick SM: Transcription factor Foxo3 controls the magnitude of T cell immune responses by modulating the function of dendritic cells. Nat Immunol 2009, 10:504–513.

60. Xie X, Lu J, Kulbokas EJ, Golub TR, Mootha V, Lindblad-Toh K, Lander ES, Kellis M: Systematic discovery of regulatory motifs in human promoters and 3’ UTRs by comparison of several mammals. Nature 2005, 434:338–345.

61. Cloonan N, Brown MK, Steptoe AL, Wani S, Chan WL, Forrest AR, Kolle G, Gabrielli B, Grimmond SM: The miR-17-5p microRNA is a key regulator of the G1/S phase cell cycle transition. Genome Biol 2008, 9:R127.

62. Trompeter HI, Abbad H, Iwaniuk KM, Hafner M, Renwick N, Tuschl T, Schira J, Muller HW, Wernet P: MicroRNAs MiR-17, MiR-20a, and MiR-106b act in concert to modulate E2F activity on cell cycle arrest during neuronal lineage differentiation of USSC. PLoS One 2011, 6:e16138.

63. Petrocca F, Visone R, Onelli MR, Shah MH, Nicoloso MS, de Martino I, Iliopoulos D, Pilozzi E, Liu CG, Negrini M, et al: E2F1-regulated microRNAs impair TGFbeta-dependent cell-cycle arrest and apoptosis in gastric cancer. Cancer Cell 2008, 13:272–286.

64. Weirauch MT, Yang A, Albu M, Cote AG, Montenegro-Montero A, Drewe P, Najafabadi HS, Lambert SA, Mann I, Cook K, et al: Determination and inference of eukaryotic transcription factor sequence specificity. Cell 2014, 158:1431–1443.

65. Chen J, Feilotter HE, Pare GC, Zhang X, Pemberton JG, Garady C, Lai D, Yang X, Tron VA: MicroRNA-193b represses cell proliferation and regulates cyclin D1 in melanoma. Am J Pathol 2010, 176:2520–2529.

66. Song B, Wang Y, Kudo K, Gavin EJ, Xi Y, Ju J: miR-192 Regulates dihydrofolate reductase and cellular proliferation through the p53-microRNA circuit. Clin Cancer Res 2008, 14:8080–8086.

67. Khella HW, Bakhet M, Allo G, Jewett MA, Girgis AH, Latif A, Girgis H, Von Both I, Bjarnason GA, Yousef GM: miR-192, miR-194 and miR-215: a convergent microRNA network suppressing tumor progression in renal cell carcinoma. Carcinogenesis 2013, 34:2231–2239.

68. Whyte WA, Orlando DA, Hnisz D, Abraham BJ, Lin CY, Kagey MH, Rahl PB, Lee TI, Young RA: Master transcription factors and mediator establish super-enhancers at key cell identity genes. Cell 2013, 153:307–319.

69. Dixon JR, Selvaraj S, Yue F, Kim A, Li Y, Shen Y, Hu M, Liu JS, Ren B: Topological domains in mammalian genomes identified by analysis of chromatin interactions. Nature 2012, 485:376–380.

70. Wickham H: Ggplot2 : elegant graphics for data analysis. New York: Springer; 2009.

71. Moreira-Filho CA, Bando SY, Bertonha FB, Silva FN, Costa Lda F, Ferreira LR, Furlanetto G, Chacur P, Zerbini MC, Carneiro-Sampaio M: Modular transcriptional repertoire and MicroRNA target analyses characterize genomic dysregulation in the thymus of Down syndrome infants. Oncotarget 2016, 7:7497–7533.

72. Reproducing iDEP analyses with auto-generated R Markdown [http://rpubs.com/ge600/R]

73. Manda K, Glasow A, Paape D, Hildebrandt G: Effects of ionizing radiation on the immune system with special emphasis on the interaction of dendritic and T cells. Front Oncol 2012, 2:102.

74. van Riggelen J, Yetil A, Felsher DW: MYC as a regulator of ribosome biogenesis and protein synthesis. Nat Rev Cancer 2010, 10:301–309.

75. Calaf GM, Hei TK: Ionizing radiation induces alterations in cellular proliferation and c-myc, c-jun and c-fos protein expression in breast epithelial cells. Int J Oncol 2004, 25:1859–1866.

76. Watson NC, Di YM, Orr MS, Fornari FA, Jr., Randolph JK, Magnet KJ, Jain PT, Gewirtz DA: Influence of ionizing radiation on proliferation, c-myc expression and the induction of apoptotic cell death in two breast tumour cell lines differing in p53 status. Int J Radiat Biol 1997, 72:547–559.

77. Qin Z, Wang PY, Su DF, Liu X: miRNA-124 in Immune System and Immune Disorders. Front Immunol 2016, 7:406.

78. Li M, Guan X, Sun Y, Mi J, Shu X, Liu F, Li C: miR-92a family and their target genes in tumorigenesis and metastasis. Exp Cell Res 2014, 323:1–6.

79. Borkowski R, Du L, Zhao Z, McMillan E, Kosti A, Yang CR, Suraokar M, Wistuba, II, Gazdar AF, Minna JD, et al: Genetic mutation of p53 and suppression of the miR-17 approximately 92 cluster are synthetic lethal in non-small cell lung cancer due to upregulation of vitamin D Signaling. Cancer Res 2015, 75:666–675.

80. Hu W, Chan CS, Wu R, Zhang C, Sun Y, Song JS, Tang LH, Levine AJ, Feng Z: Negative regulation of tumor suppressor p53 by microRNA miR-504. Mol Cell 2010, 38:689–699.

81. Bublik DR, Bursac S, Sheffer M, Orsolic I, Shalit T, Tarcic O, Kotler E, Mouhadeb O, Hoffman Y, Fuchs G, et al: Regulatory module involving FGF13, miR-504, and p53 regulates ribosomal biogenesis and supports cancer cell survival. Proc Natl Acad Sci U S A 2017, 114:E496–E505.

82. Parveen A, Akash MS, Rehman K, Kyunn WW: Dual Role of p21 in the Progression of Cancer and Its Treatment. Crit Rev Eukaryot Gene Expr 2016, 26:49–62.

83. Gallant S, Gilkeson G: ETS transcription factors and regulation of immunity. Arch Immunol Ther Exp (Warsz) 2006, 54:149–163.

84. Fendler W, Malachowska B, Meghani K, Konstantinopoulos PA, Guha C, Singh VK, Chowdhury D: Evolutionarily conserved serum microRNAs predict radiation-induced fatality in nonhuman primates. Sci Transl Med 2017, 9.

85. Park D, Kim H, Kim Y, Jeoung D: miR-30a Regulates the Expression of CAGE and p53 and Regulates the Response to Anti-Cancer Drugs. Mol Cells 2016, 39:299–309.

86. Ioannidis JP: Why most published research findings are false. PLoS Med 2005, 2:e124.

87. Trapnell C, Roberts A, Goff L, Pertea G, Kim D, Kelley DR, Pimentel H, Salzberg SL, Rinn JL, Pachter L: Differential gene and transcript expression analysis of RNA-seq experiments with TopHat and Cufflinks. Nat Protoc 2012, 7:562–578.

88. Patel AP, Tirosh I, Trombetta JJ, Shalek AK, Gillespie SM, Wakimoto H, Cahill DP, Nahed BV, Curry WT, Martuza RL, et al: Single-cell RNA-seq highlights intratumoral heterogeneity in primary glioblastoma. Science 2014, 344:1396–1401.

89. Liu ZP, Wu C, Miao H, Wu H: RegNetwork: an integrated database of transcriptional and post-transcriptional regulatory networks in human and mouse. Database (Oxford) 2015, 2015.

90. Friard O, Re A, Taverna D, De Bortoli M, Cora D: CircuitsDB: a database of mixed microRNA/transcription factor feed-forward regulatory circuits in human and mouse. BMC Bioinformatics 2010, 11:435.

91. Jiang C, Xuan Z, Zhao F, Zhang MQ: TRED: a transcriptional regulatory element database, new entries and other development. Nucleic Acids Res 2007, 35:D137–140.

92. Zheng G, Tu K, Yang Q, Xiong Y, Wei C, Xie L, Zhu Y, Li Y: ITFP: an integrated platform of mammalian transcription factors. Bioinformatics 2008, 24:2416–2417.

93. Neph S, Stergachis AB, Reynolds A, Sandstrom R, Borenstein E, Stamatoyannopoulos JA: Circuitry and dynamics of human transcription factor regulatory networks. Cell 2012, 150:1274–1286.

94. Marbach D, Lamparter D, Quon G, Kellis M, Kutalik Z, Bergmann S: Tissue-specific regulatory circuits reveal variable modular perturbations across complex diseases. Nat Methods 2016, 13:366–370.

95. Consortium EP: An integrated encyclopedia of DNA elements in the human genome. Nature 2012, 489:57–74.

96. Han H, Shim H, Shin D, Shim JE, Ko Y, Shin J, Kim H, Cho A, Kim E, Lee T, et al: TRRUST: a reference database of human transcriptional regulatory interactions. Sci Rep 2015, 5:11432.

97. Wong N, Wang X: miRDB: an online resource for microRNA target prediction and functional annotations. Nucleic Acids Res 2015, 43:D146–152.

98. Agarwal V, Bell GW, Nam JW, Bartel DP: Predicting effective microRNA target sites in mammalian mRNAs. Elife 2015, 4.

99. Chou CH, Shrestha S, Yang CD, Chang NW, Lin YL, Liao KW, Huang WC, Sun TH, Tu SJ, Lee WH, et al: miRTarBase update 2018: a resource for experimentally validated microRNA-target interactions. Nucleic Acids Res 2018, 46:D296-D302.

